# Genome-wide association studies for yield-related traits in soft red winter wheat grown in Virginia

**DOI:** 10.1101/471656

**Authors:** Brian P. Ward, Gina Brown-Guedira, Frederic L. Kolb, David A. Van Sanford, Priyanka Tyagi, Clay H. Sneller, Carl A. Griffey

## Abstract

Grain yield is a trait of paramount importance in the breeding of all cereals. In wheat (*Triticum aestivum* L.), yield has steadily increased since the Green Revolution, though the current rate of increase is not forecasted to keep pace with demand due to growing world population and affluence. While several genome-wide association studies (GWAS) on yield and related component traits have been performed in wheat, the previous lack of a reference genome has made comparisons between studies difficult. In this study, a GWAS for yield and yield-related traits was carried out on a population of 324 soft red winter wheat lines across a total of four rain-fed environments in the state of Virginia using single-nucleotide polymorphism (SNP) marker data generated by a genotyping-by-sequencing (GBS) protocol. Two separate mixed linear models were used to identify significant marker-trait associations (MTAs). The first was a single-locus model utilizing a leave-one-chromosome-out approach to estimating kinship. The second was a sub-setting kinship multi-locus method (FarmCPU). The single-locus model identified nine significant MTAs for various yield-related traits, while the FarmCPU model identified 74 significant MTAs. The availability of the wheat reference genome allowed for the description of MTAs in terms of both genetic and physical positions, and enabled more extensive post-GWAS characterization of significant MTAs. The results indicate promising avenues for increasing grain yield by exploiting variation in traits relating to the number of grains per unit area, as well as phenological traits influencing grain-filling duration of genotypes.

## Introduction

Worldwide, wheat has the fourth-highest production of all crops, with a net production value that is second-highest of any crop [1]. In addition, wheat maintains the highest global harvested acreage of any crop [2]. To keep pace with an increasing world population and changes in diets due to increasing affluence, worldwide cereal production will have to increase by an estimated 50% over the period ending in 2050, requiring continuing genetic gains in yield potential of approximately 1.1% per year [3]. Sharma, et al. [4] estimated a historical average increase in grain yields in spring wheat of 0.65% per year when analyzing data across 15 years and 919 environments, while a recent study involving winter wheat in the Eastern United States estimated yearly increases in grain yield between 0.56% and 1.41%, depending upon environment [5].

Improvements to yield via direct selection are hampered by its highly quantitative and polygenic nature. Selection based on different yield component traits and crop physiology theory may offer additional avenues for increasing genetic gain in yield while avoiding yield plateaus [6]. However, relationships between traits must be taken into account, as there are many negative correlations between related traits, such as the well-documented correlation between the number of grains m^-2^ and average grain size [7]. A wide body of literature suggests that wheat is primarily sink-limited with respect to production of photosynthetic assimilates (reviewed in [8]). Supporting this theory, several studies have suggested that maximizing the number of seeds per unit area is critical for avoiding assimilate sink limitations and maximizing yield (reviewed in [9]). The two possible routes for increasing seeds per unit area are to: 1) increase the average number of seeds per head, and/or 2) increase the number of heads per unit area. Increases in yield will require both the production of greater numbers of grains, and an increase in the availability of photosynthates to prevent a corresponding drop in kernel sizes due to the compensation between these two traits [10]. Several previous studies have focused on increasing photosynthate availability via alterations to canopy light interception. Reynolds et al. [9] note that the amount of light interception is already nearly maximized in many wheat cultivars during the period after canopy closure and prior to leaf senescence. This leaves traits increasing the duration of light interception, e.g. faster canopy establishment through increased early-season vigor or later senescence via the “stay-green” trait [11], as appealing avenues for increasing photosynthate production.

In winter wheat, several genes are known to exert major effects on traits affecting seed production per unit area and photosynthesis duration. The time required for plants to flower and ultimately reach maturation is affected by multiple homeologous copies of the *VRN* vernalization requirement genes and the *PPD* photoperiod response genes [12]. Multiple major-effect quantitative trait loci (QTLs) affecting grain weight have also been identified, including *TaSus2-2B* [13], the *TaGW2* homeologous genes [14], and cell wall invertase (*Cwi*) genes [15]. However, it is likely that many genes affecting yield-related traits have yet to be identified. Genome-wide association studies (GWAS) and linkage mapping are the two predominate methods employed in plant breeding for associating phenotypic variation with underlying genetic variation. GWAS offers higher resolution due to many more ancestral gene recombinations within the testing panel, as opposed to only one or a few meiotic recombinations in a linkage mapping population. However, allele frequency is a primary factor limiting power in GWAS studies; marker-trait associations (MTAs) will be difficult or impossible to detect if causal variants are rare within the testing population [16]. Relatively few GWAS analyses for yield and yield-related traits have been conducted in wheat, and of these, fewer still have been conducted in winter wheat germplasm [17–20]. Association studies have been more common in spring wheat, though the majority of these have been candidate-gene studies, with genome-wide studies being much more limited [21]. Finally, it has been more common to perform GWAS in crop species using assembled diversity panels, rather than elite germplasm in current use by breeding programs [22]. GWAS in elite germplasm is typically limited to the identification of smaller-effect MTAs, as MTAs of major effect will have likely already become fixed within the mapping population [23]. Nevertheless, GWAS using panels of elite germplasm remain useful due to their higher relevance to the process of cultivar development [22]. One limitation of previous wheat GWAS studies was the lack of a reference genome, making comparisons between studies’ findings difficult. The recent publication of a full chromosome-anchored genome assembly [24] should allow for better curation of MTAs uncovered by GWAS and linkage mapping studies.

The present study sought to perform GWAS in a panel of elite soft winter wheat lines sourced from breeding programs in the Eastern and Midwestern United States. Genotyping was performed via genotyping-by-sequencing (GBS), using genetic maps and anchoring SNPs to the v1.0 Chinese Spring wheat reference genome, allowing for determination of both physical and genetic positions of identified MTAs.

## Materials and methods

### Germplasm selection

The study was conducted over two years, and included a total of 329 genotypes (**S1 Table**). Of these, 41 genotypes were tested in both years. Of the remaining genotypes, half (144) were tested only in the first year, and the other half (144) were tested only in the second year. Within each year, genotypes were sourced from breeding programs in Illinois (31), Kentucky (30), Missouri (2), and Virginia (122). Seven genotypes were removed during quality filtering of the genotypic data, leaving a total of 322 genotypes among both years used for further analysis. Five checks were included in the study, including ‘Bess’, ‘Branson’, IL00-8530, ‘Roane’, and ‘Shirley’. These lines were selected as they are soft wheat lines that have frequently been grown in production and used as parents in the mid-Atlantic and Midwest regions. With the exception of checks and several older cultivars, the majority of genotypes were either F_4_ or F_5_ filial generation.

## Phenotyping

### Experimental design and field management

Experimental plots were planted in a total of four environments (two locations in two years) in the 2013-14 and 2014-15 winter wheat growing seasons. Within each year, trials were planted at Kentland Farm near Blacksburg, Virginia (Guernsey/Hayter silt loams, 37.1965° N, 80.5718° W, 531 m elevation) and the Eastern Virginia Agricultural Research and Extension Center in Warsaw, Virginia (Kempsville sandy loam, 37.9879° N, 76.7770° W, 40 m elevation). A generalized randomized complete block design (GRCBD) with two replications was used in each environment.

Each experimental unit consisted of a seven-row plot with a length of 2.74 m, width of 0.91 m, row-spacing of 15.2 cm, and a harvested area of 2.49 m^2^. All plots were sown with 70 g of seed. Seed was treated with Raxil^®^ MD fungicide (0.48% tebuconazole/0.64% metalaxyl; Bayer CropScience) at a rate of 2.95 mL a.i. per kg of seed, and Gaucho^®^ 600 flowable insecticide (48.7% imidacloprid; Bayer CropScience) at a rate of 0.7 mL a.i. per kg of seed. At each location, seeds were planted to roughly coincide with the average date of first frost.

At Blacksburg and Warsaw, several tiller counts representative of the test area as a whole were used to calculate ideal nitrogen application rates at Zadok’s growth stage 25 [25] in the spring. Once plants reached Zadok’s growth stage 30, tissue samples were collected by cutting handfuls of leaf material from roughly 25 randomly-selected sites per environment. For each environment, collected leaf tissue was then mixed thoroughly and analyzed for nitrogen content by the Dumas method (reviewed in [26]) to determine ideal additional nitrogen application rates, per standard regional recommendations from the Virginia Cooperative Extension Service [27]. Chemicals were applied as needed to control lodging and pest pressure in each environment.

### Data collection

**S2 Table** summarizes the phenotypic traits that were assessed across all environments, with their abbreviations, units of measure, and trait ontologies. Normalized Difference Vegetative Index (NDVI) was measured for each plot at Zadok’s growth stage 25 as described by Phillips et al. [28] using a Greenseeker^®^ Handheld crop sensor (Trimble^®^ Agriculture, Sunnyvale, CA). Heading date (HD) was recorded as the Julian date at which 50% of plant tillers within a plot had extruded heads from the boot. Physiological maturity date (MAT) was defined as the date at which 50% of peduncles within a plot had turned yellow. After plants reached maturity, a 0.914 m cutting of all above-ground plant material was taken from one of the three inner rows of each plot and placed inside a paper bag. The number of spikes per cutting were counted manually to derive an estimate of spikes per square meter (SSQM; count). Cuttings were then threshed on a plot combine (Wintersteiger NA Inc., Salt Lake City, UT) with settings optimized to recover as much threshed seed as possible. Threshed seeds were weighed to derive an estimate of grain weight per meter of row (GW; grams). The total number of seeds threshed from each cutting were then counted on a Count-A-Pak optical seed counter (Seedburo^®^ Equipment, Des Plaines, IL) to derive an estimate of grains per square meter (GSQM; count). The number of heads was divided by the number of seeds to derive an estimate of seeds per head (SPH; count). Thousand-kernel weight (TKW; grams) was then calculated as the total number of seeds threshed from the cutting divided by their weight multiplied by 1,000. Plant height (HT; cm) was averaged from two measurements within each plot, and was recorded as the distance from the soil surface to the tip of the heads (excluding any awns if present). Plots were harvested at maturity using a Wintersteiger plot combine. Moisture content and test weight of harvested grain was measured using a GAC^®^ 2500-AGRI grain analysis computer (Dickey-John^®^ Corporation, Auburn, IL). Grain yield (YLD; kg ha^-1^) was derived from the yield of each harvested plot at 13.5% moisture equivalence.

Whole grain starch (STARCH; percent), and protein (PROT; percent) content were estimated via near-infrared (NIR) spectroscopy for subsamples from each plot using an XDS Rapid Content Analyzer (FOSS NIR Systems, Laurel, MD). Thirty samples, each consisting of 25 grams of seed, were sent to Cumberland Valley Analytical Services (Hagerstown, MD) for wet-chemistry analysis of protein content [29] to generate calibration curves for the NIR data. Samples were chosen to represent the full range of observed NIR protein content values. The coefficient of determination of a regression between NIR and wet-chemistry derived protein content values was 0.78 (data not shown).

### Spatial corrections

For each individual trait/environment combination, an *ad-hoc* correction for field heterogeneity was performed using two-dimensional tensor product penalized B-splines [30], implemented in a mixed model framework in the R [31] package ‘SpATS’ [32]. The package default parameters were used to fit cubic splines with quadratic penalization functions in both row and column dimensions. For a particular environment with plots arranged in *m* rows and *n* columns, the number of knots used to fit splines was set to (⌊*m*⌋/2) − 1 and (⌊*n*⌋/2) − 1, respectively. Fitted values generated by the SpATS model were used for subsequent modeling of phenotypes across environments, as described in the section below.

Heritability for each trait/environment combination was estimated using the method of Cullis, et al. [33]:

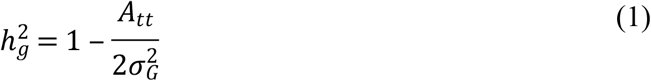

Where the generalized heritability 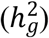 is a function of the average pairwise prediction error between pairs of genotypes within an environment (*A_tt_*), and the genotypic variance 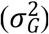. Heritability estimates were calculated from the models fit by the SpATS package, and were compared against the heritability estimates generated by a baseline model:

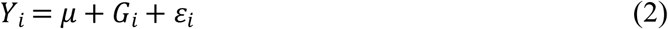

Where phenotypic response (*Y_i_*) is a function of the within-environment mean (μ), the fixed effect of the *i*th genotype (*G_i_*), and residual error (*ε_i_*).

### Modelling of phenotypes across environments

Each location/year combination was considered as a unique environment in order to model phenotypic response across environments. For each trait, the following random effects ANOVA model was fit using the *lme4* package [34] in the R statistical computing environment [31]:

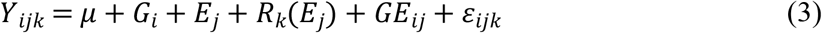

Where the phenotypic response (*Y_ijk_*) is a function of the overall mean (*μ*), the ith genotype (*G_i_*), the kth replication (*R_k_*) nested within the jth environment (*E_j_*), the genotype-environment interaction (*GE_ij_* and the residual error (*ε_ijk_*). Genotypic best-linear unbiased predictors (BLUPs) were calculated for use as the phenotypic input for the subsequent GWAS analyses.

### Genotyping

Genomic DNA was isolated from fresh green seedling leaf tissue using an LGC Genomics Oktopure^™^ robotic extraction platform with sbeadex^™^ magnetic microparticle reagent kits. Genotyping-by-sequencing was performed using an Illumina HiSeq 2500 following a double digest of genomic DNA using the restriction enzymes *PstI* and *MseI*, as described in Poland et al., 2012 [35]. SNP calling was performed using TASSEL-GBS v5.2.43 [36,37]. The Burrows-Wheeler Aligner [38] v0.7.17-r1188 was used to align Illumina-generated short reads to the International Wheat Genome Sequencing Consortium (IWGSC) v1.0 wheat reference sequence [24]. The raw genotypic data was filtered to retain only biallelic SNPs, and to remove SNPs with missing data frequencies > 50%, mean sequencing depth < 2, heterozygous call frequencies > 15%, or minor allele frequency < 5%. SNPs that aligned to unmapped contigs were removed. Missing data was then imputed using Beagle v4.1 with default settings [39]. After imputation, the genotypic data was filtered a second time to remove SNPs with minor allele frequency < 5% or heterozygous call frequency < 15%. PLINK 1.9 [40] was used to remove all but one SNP in groups of SNPs in perfect linkage disequilibrium (LD; r^2^ = 1). After filtering, 29,949 SNPs remained for further analysis.

### Assays for polymorphisms of major effect

Several genes and polymorphisms of major effect were assayed using LGC Genomics KASPar SNP assays. All included assays are listed with primer sequences in **S3 Table-A**. Briefly, they included assays for the 1RS:1AL and 1RS:1BL translocations from rye (*Secale cereale*), polymorphisms within the *Ppd-A1, Ppd-B1*, and *Ppd-D1* photoperiod sensitivity genes located on chromosomes 2A, 2B, and 2B respectively, the *Rht-B1* and *Rht-D1* dwarfing genes located on chromosomes 4B and 4D respectively, and polymorphisms of the *Vrn-A1* and *Vrn-B1* genes located on chromosomes 5A and 5B.

KASP assays for polymorphisms within exon 4 and exon 7 of *Vrn-A1* were included in this study. The molecular mechanism behind *Vrn-A1*’s effects on vernalization requirement remain contested. Chen et al. [41] proposed that a SNP occurring in exon 4 of *Vrn-A1* was the causal locus differentiating between the *Vrn-A1a* short vernalization requirement allele present in the cultivar Jagger and the *Vrn-A1b* long vernalization requirement allele present in the cultivar ‘2174’. Diaz et al. [42] later proposed that the differences between *Vrn-A1a* and *Vrn-A1b* vernalization requirements are due to *Vrn-A1* copy number variations. Li et al. [43] subsequently reported that although Jagger contains two copies of *Vrn-A1*, whereas ‘2174’ contains three copies of the gene, differences in vernalization requirements between the two cultivars are in fact due to structural differences, particularly due to a SNP located in exon 7, converting the 180^th^ amino acid residue from alanine in *Vrn-A1a* to valine in *Vrn-A1b*. Finally, Kippes et al. [44] proposed that the effects of *Vrn-A1* arise from the product of gene *VRN-D4* disrupting the binding of the RNA-binding repressor *TaGRP2* to *Vrn-A1*. The KASP assays for *Vrn-A1* used in this study have been shown to be suggestive, but not perfect predictors of vernalization requirement due to *Vrn-A1* alleles.

In addition, assays for the *Sr36* stem rust (*Puccinia graminis* Pers.) resistance gene, and the sucrose-synthase gene *TaSus2-2B*, which affects kernel weight, were included. Gene *Sr36* is located on a 2G:2B alien translocation originating from *Triticum timopheevi* [Zhuk.] Zhuk. (A^m^A^m^GG) [45,46]. Gene *TaSus2-2B* is located on the short arm of chromosome 2B, and is one of the three sucrose synthase *Sus2* orthologs located on chromosomes 2A, 2B, and 2D [13]. Two common haplotypes for *TaSus2-2B* include *Hap-H* (high seed weight) residing on the 2G:2B translocation, and *Hap-L* (low seed weight). Cabrera et al. [47] found that the simple sequence repeat (SSR) marker *Xwmc477* used to test for the 2G:2B translocation was in perfect LD with *TaSus2-2B* alleles, suggesting that the haplotypes of this gene can also be used to test for the presence or absence of the 2G:2B translocation.

### Population structure and linkage disequilibrium

Prior to performing GWAS, population structure was examined via principle component analysis (PCA) of the filtered and imputed genotypic data using the SNPRelate package [48] in R. Linkage disequilibrium was calculated on a pairwise basis for 4,500 SNPs randomly sampled from each genome, yielding 173,891 comparisons. LD decay was plotted for the A, B, and D genomes separately by randomly selecting 20,000 of the pairwise LD comparisons from each genome. LD was likewise plotted for each separate chromosome using 20,000 randomly-selected pairwise comparisons from each chromosome. In addition, 500 randomly selected SNPs were used to estimate inter-chromosomal LD, yielding a total of 118,175 comparisons. For intra-chromosomal SNPs, r^2^ values for pairwise LD comparisons were plotted against physical distance, and a second-degree locally-weighted scatterplot smoothing (LOESS) curve was fit to the data [49]. The 98^th^ percentile of the LD distribution for non-linked (i.e. inter-chromosomal) loci was defined as the linkage-disequilibrium critical value; all r^2^ values exceeding this value for intra-chromosomal loci were assumed to have been caused by genetic linkage [17]. In addition, LD was calculated between the KASP *Sr36* assay and all other SNPs present on chromosome 2B. Finally, the fixation index (*F_ST_*) was calculated for each SNP using Weir and Cockerham’s estimator [50]. All LD and *F_ST_*-related calculations were performed using VCFtools v0.1.17 [51].

### Genome-wide association analysis

For each trait, single-locus mixed linear model genome-wide association analyses were performed with the Genome-Wide Complex Trait Analysis (GCTA) software [52], using a leave-one-chromosome-out (LOCO) method in which a separate genetic relationship matrix (GRM) is estimated from SNP data for each chromosome. Specifically, the LOCO approach entails excluding all SNPs located on the chromosome of the SNP undergoing testing when estimating the GRM. Adjusted p-values for each SNP were calculated using the method of Benjamini and Hochberg [53], and SNPs with adjusted p-values below 0.05 were considered significant.

In addition, GWAS was performed for each trait using the Fixed and Random Model Circulating Probability Unification (FarmCPU) model [54], using the R package ‘FarmCPUpp’ [55]. The same significance threshold applied in the single-locus tests was used for the FarmCPU results. To enhance our confidence in significant MTAs identified by FarmCPU, we implemented a bootstrapping method utilized by Wallace, et al. [56], in which 10% of the phenotypic observations were randomly replaced with missing data for a total of 100 runs of the model. Subsequently, for each trait the resample model inclusion probability [57] was calculated for each SNP by determining the fraction of bootstraps in which its adjusted p-value exceeded the significance threshold. The value 0.1 was chosen as a lower threshold for the RMIP as it coincided with the point of inflection in the RMIP density curves (data not shown). For each model, the first four principal components were included to model population structure, based upon visual examination of the Scree plot for variance explained by each PC, and clustering of lines shown in biplots of the first few PCs (**S1 Fig**.).

### Candidate genes and translation effects

Haplotype blocks surrounding each significant MTA were identified using the method of Gabriel et al. [58] by running the --blocks command in PLINK 1.9. Some significant MTAs did not reside within any larger haplotype block, while some haplotype blocks contained multiple significant MTAs. Subsequently, all genes overlapping significant MTAs and associated haplotype blocks were identified using the IWGSC v1.1 RefSeq annotation [24] with functional annotations from the IWGSC v1.0 annotation. In addition, Ensembl identifiers were retrieved for all genes overlapping with significant SNPs or haplotype blocks. A table of wheat genes with trEMBL or Swissprot-generated protein annotations in UniProt was downloaded and used to identify all wheat genes with predicted functions located within 2Mb of each significant MTA. The Ensembl Variant Effect Predictor [59] was then used to classify significant SNPs as being either intergenic, intronic, exonic, or upstream/downstream proximal variants. The predicted allele substitution effects of exonic SNPs on protein translation were classified as synonymous, missense, or nonsense. For gene-proximal SNPs, the distance to the closest gene was recorded.

## Results

### Trait heritability and correlations

The use of spatially-corrected data tended to drastically increase generalized heritability within each environment. The trait GW had the lowest mean generalized heritability, averaging 0.2 across environments for raw, uncorrected data, and 0.33 after performing spatial corrections (**Table 1**). The trait TKW had the highest mean generalized heritability when using uncorrected data 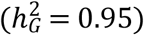, though heritability was slightly decreased when using spatially-corrected data 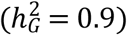. When spatially-corrected data was used, TWT had the highest average heritability 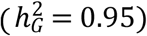. YLD had a moderate mean, across-environment heritability of 0.57 when calculated using uncorrected data, though this increased to 0.81 when using spatially-corrected data. Pearson correlation coefficients were calculated for each pair of traits using the phenotypic BLUPs (**Table 2**). Phenological traits (HD, FLSG, and MAT) all displayed a high degree of intercorrelation. This was also the case for traits relating to grain size and density per unit area (TKW, GSQM, SSQM, and SPH). For instance, the traits GSQM and TKW demonstrated a strong negative correlation, as predicted due to trait compensation effects. Critically, only weak correlation was observed between traits relating to grain size/density and phenological traits, suggesting that these two classes of traits could be improved independently. YLD was most highly correlated with the traits GW and MAT (positive), and grain protein (negative). The negative correlation between yield and grain protein content has been well documented in the past (e.g. [60–62]).

**Table 1:** Trait descriptive statistics and generalized heritability estimates calculated using raw plot values and spatially-adjusted values.

| Trait <sup>a</sup> | Units | Descriptive Statistics |  |  |  | Generalized Heritability |  |  |  |  |  |  |  |  |  |
| --- | --- | --- | --- | --- | --- | --- | --- | --- | --- | --- | --- | --- | --- | --- | --- |
|  |  | min | mean | max | SD | Raw Plot Values |  |  |  |  | Spatially-Adjusted Values |  |  |  |  |
|  |  |  |  |  |  | 14Bb | 14War | 15Bb | 15War | mean | 14Bb | 14War | 15Bb | 15War | mean |
| FLSG | days | 21 | 28.69 | 38 | 2.76 | 0.26 | 0.8 | 0.75 | 0.55 | 0.59 | 0.55 | 0.83 | 0.76 | 0.75 | 0.72 |
| GSQM | Grains m-2 | 8460 | 1.85E+04 | 3.13E+04 | 3277 | 0.62 | 0.5 | 0.51 | 0.58 | 0.55 | 0.71 | 0.51 | 0.54 | 0.67 | 0.61 |
| GW | g dwt m-1 row | 47.68 | 96.64 | 157.9 | 16.53 | 0.19 | 0.14 | 0.44 | 0.05 | 0.2 | 0.43 | 0.16 | 0.47 | 0.25 | 0.33 |
| HD | Julian days (Jan1) | 121 | 128 | 136 | 3.21 | 0.75 | 0.94 | 0.95 | 0.96 | 0.9 | 0.82 | 0.95 | 0.96 | 0.97 | 0.92 |
| HT | cm | 59.69 | 85.43 | 119.4 | 9.29 | 0.85 | 0.85 | 0.84 | 0.86 | 0.85 | 0.92 | 0.88 | 0.88 | 0.87 | 0.89 |
| MAT | Julian days (Jan1) | 151 | 159.3 | 171 | 4.72 | 0.83 | 0.87 | 0.86 | 0.76 | 0.83 | 0.89 | 0.91 | 0.89 | 0.9 | 0.9 |
| NDVI | - | 0.26 | 0.54 | 0.75 | 0.08 | 0.06 | 0.39 | 0.41 | 0.41 | 0.32 | 0.7 | 0.75 | 0.68 | 0.64 | 0.69 |
| PROT | % | 9.67 | 12.34 | 16.04 | 1.01 | 0.66 | 0.38 | 0.63 | 0.74 | 0.6 | 0.67 | 0.4 | 0.71 | 0.78 | 0.64 |
| SPH | count | 8.54 | 21.94 | 33.29 | 3.04 | 0.86 | 0.82 | 0.77 | 0.81 | 0.82 | 0.85 | 0.82 | 0.75 | 0.84 | 0.82 |
| SSQM | Spikes m-2 | 459.3 | 853 | 1485 | 161.2 | 0.6 | 0.62 | 0.45 | 0.52 | 0.55 | 0.65 | 0.63 | 0.5 | 0.63 | 0.6 |
| STARCH | % | 46.88 | 52.51 | 56.49 | 1.41 | 0.46 | 0.71 | 0.19 | 0.69 | 0.51 | 0.8 | 0.72 | 0.73 | 0.83 | 0.77 |
| TKW | grams | 24.1 | 34.57 | 91.6 | 3.98 | 0.96 | 0.97 | 0.94 | 0.93 | 0.95 | 0.97 | 0.98 | 0.69 | 0.97 | 0.9 |
| TWT | g L-1 | 652.6 | 759 | 810.9 | 19.7 | 0.96 | 0.92 | 0.91 | 0.93 | 0.93 | 0.98 | 0.96 | 0.9 | 0.95 | 0.95 |
| YLD | kg ha-1 | 3579 | 6627 | 9053 | 1027 | 0.63 | 0.45 | 0.73 | 0.46 | 0.57 | 0.84 | 0.81 | 0.78 | 0.81 | 0.81 |

14Bb Blacksburg, VA 2014; 14War Warsaw, VA 2014; 15Bb Blacksburg, VA 2015; 15War Warsaw, VA 2015
<sup>a</sup> FLSG flag leaf stay green; GSQM grains per square meter; GW grain weight; HD heading date; HT plant height; MAT physiological maturity date; NDVI normalized-difference vegetation index at Zadok's GS25; PROT wet chemistry-validated whole-grain protein content; SPH seeds per head; SSQM spikes per square meter; STARCH whole-grain starch content; TKW thousand kernel weight; TWT test weight; YLD grain yield

**Table 2:** Phenotypic correlations among traits calculated using the across-environment genotype BLUPs.

|  | FLSG | GSQM | GW | HD | HT | MAT | NDVI | SPH | SSQM | STARCH | TKW | TWT | PROT | YLD |
| --- | --- | --- | --- | --- | --- | --- | --- | --- | --- | --- | --- | --- | --- | --- |
| FLSG | 1 |  |  |  |  |  |  |  |  |  |  |  |  |  |
| GSQM | 0.01 | 1 |  |  |  |  |  |  |  |  |  |  |  |  |
| GW | 0.27 | 0.64 | 1 |  |  |  |  |  |  |  |  |  |  |  |
| HD | -0.43 | 0.14 | -0.06 | 1 |  |  |  |  |  |  |  |  |  |  |
| HT | -0.18 | -0.16 | -0.21 | 0.15 | 1 |  |  |  |  |  |  |  |  |  |
| MAT | 0.18 | 0.26 | 0.19 | 0.71 | -0.07 | 1 |  |  |  |  |  |  |  |  |
| NDVI | 0.05 | 0.05 | 0.08 | 0.12 | 0.13 | 0.1 | 1 |  |  |  |  |  |  |  |
| SPH | 0 | 0.48 | 0.4 | 0.12 | 0.02 | 0.21 | -0.2 | 1 |  |  |  |  |  |  |
| SSQM | 0.02 | 0.53 | 0.24 | 0.02 | -0.18 | 0.05 | 0.25 | -0.48 | 1 |  |  |  |  |  |
| STARCH | 0.2 | 0.3 | 0.32 | -0.04 | -0.23 | 0.19 | -0.12 | 0.29 | 0.01 | 1 |  |  |  |  |
| TKW | 0.25 | -0.66 | 0.14 | -0.24 | -0.01 | -0.13 | 0.01 | -0.24 | -0.43 | -0.07 | 1 |  |  |  |
| TWT | 0.04 | -0.22 | -0.19 | -0.28 | 0.18 | -0.31 | -0.08 | -0.2 | -0.02 | -0.25 | 0.1 | 1 |  |  |
| PROT | 0.07 | -0.41 | -0.39 | -0.15 | 0.11 | -0.21 | 0.15 | -0.39 | -0.01 | -0.62 | 0.15 | 0.25 | 1 |  |
| YLD | 0.32 | 0.36 | 0.58 | 0.18 | -0.08 | 0.48 | 0.13 | 0.33 | 0.03 | 0.41 | 0.11 | -0.2 | -0.45 | 1 |

FLSG flag leaf stay green; GSQM grains per square meter; GW grain weight; HD heading date; HT plant height; MAT physiological maturity date; NDVI normalized-difference vegetation index at Zadok's GS25; PROT wet chemistry-validated whole-grain protein content; SPH seeds per head; SSQM spikes per square meter; STARCH whole-grain starch content; TKW thousand kernel weight; TWT test weight; YLD grain yield

### Polymorphisms of major effect

An estimation of allele effects for the KASP markers assaying genes of known function revealed that the stem rust resistance gene *Sr36* and the sucrose-synthase gene *TaSus2-2B* produced many significant differences among genotypes for multiple traits (**S3 Table-B**). These two genes also consistently produced significant differences of similar magnitudes for the same traits. The presence of the *Rht-B1b* and *Rht-D1b* dwarfing alleles produced significant effects of opposite signs for many traits, including SSQM, TKW, and TWT. Effects were also of opposite signs for HT and YLD, though in both of these cases the effects of *Rht-B1b* were not significant. In general, many of the genes of known function did not produce significant effects within the testing panel. For instance, of the three *Ppd* genes, only one (*Ppd-D1*) had a significant effect on HD, while only *Ppd-B1* generated significant effects on the phenological traits of FLSG and MAT. The effects of *Vrn* vernalization genes on phenological traits were likewise sporadic, with only the *Vrn-B1* polymorphism exhibiting a significant effect on FLSG, and only the *Vrn-A1* exon 4 polymorphism exhibiting significant effects on HD and MAT. In addition, none of the loci of major effect assayed with KASP markers were classified as significant in the GWAS. Some potential reasons for these findings will be discussed below.

### Population structure and linkage disequilibrium

Principle component analysis of the processed GBS SNP data revealed substantial admixture among genotypes, with the first principle component only explaining 7.84% of the total genotypic variance (**S1 Fig**). Lines included in the panel formed two distinct clusters in the biplot of the first two principle components. These two clusters were largely delineated by the presence or absence of the 2G:2B translocation as determined by the *Sr36* KASP assay (**Fig 1-A**; similar results for *TaSus2-2B* not shown). In contrast, neither the 1BL:1RS nor the 1AL:1RS alien translocations from rye (*Secale cereal* L.) produced any discernable clustering of genotypes (data not shown). When the SNP data was thinned to remove SNPs in high LD with each other (i.e. limiting the maximum pairwise LD between SNPs to r^2^ = 0.2), the population stratifying effects of *Sr36* and the underlying 2G:2B translocation were removed, and the first principle component explained 3.68% of variation (**Fig 1-B**).

**Fig 1.**
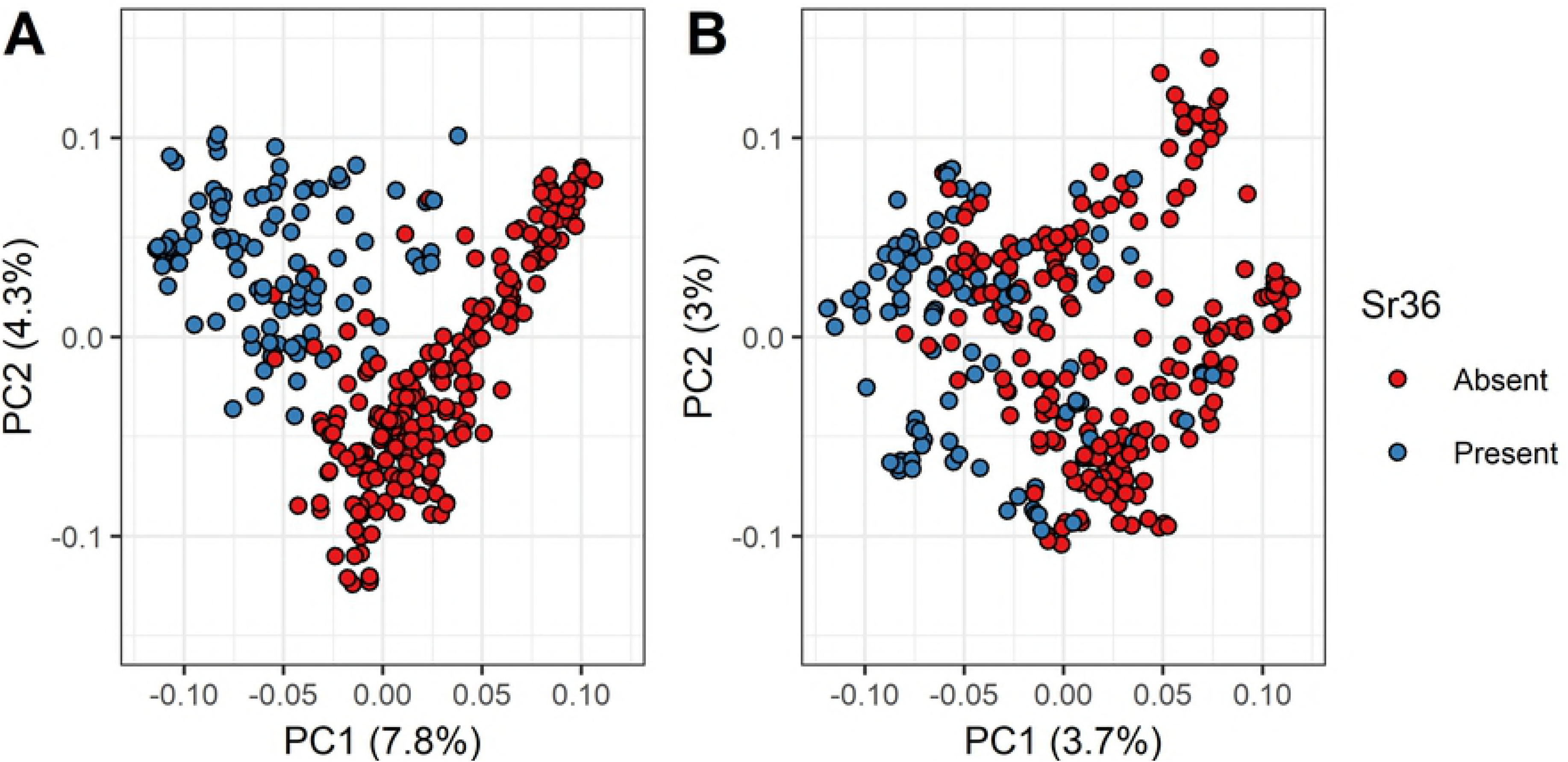
Biplots of Genotypic Principal Components. The first and second principal components of the SNP matrix were plotted against each other, using (A) all SNPs that passed initial filtering parameters, and (B) a thinned subset of SNPs selected to be in approximate linkage equilibrium, so that no pair of SNPs displayed significant LD (r^2^ > 0.2). The percent of the total genotypic variance explained is listed on each axis. Genotypes are divided into two groups based on the presence of absence of the *Sr36* stem rust resistance gene and underlying 2G:2B translocation as determined by KASP assay.

Linkage disequilibrium decay plots demonstrated significant LD extending out to large physical distances. Notably, intra-chromosomal LD in the B genome extended much farther than in either the A or D genomes (**Fig. 2**). Within the B genome, chromosome 2B displayed the most extensive LD, chromosomes 4B and 5B displayed intermediate LD, and chromosomes 1B, 3B, 6B, and 7B displayed low LD more similar to the overall LD patterns observed in the A and D genomes. The genome-wide *F_ST_* scan following the splitting of the overall panel based on the result of the *Sr36* KASP assay demonstrated that the 2G:2B translocation formed a large block of high-LD SNPs spanning almost all of chromosome 2B (**Fig. 3**). In addition, an enrichment of high-*F_ST_* SNPs on chromosomes 2A and 2D suggested misalignment of SNPs among the group 2 homeologous chromosomes. Further examination of *F_ST_* values on chromosome 2B indicated a general linear relationship between *F_ST_* and the value of r^2^ measured against the *Sr36* KASP marker for most SNPs, with a minority of SNPs exhibiting high *F_ST_* with little or no corresponding LD with the *Sr36* KASP marker (**Fig. 4-A**). These SNPs were distributed throughout the chromosome (**Fig. 4-B**).

**Fig 2.**
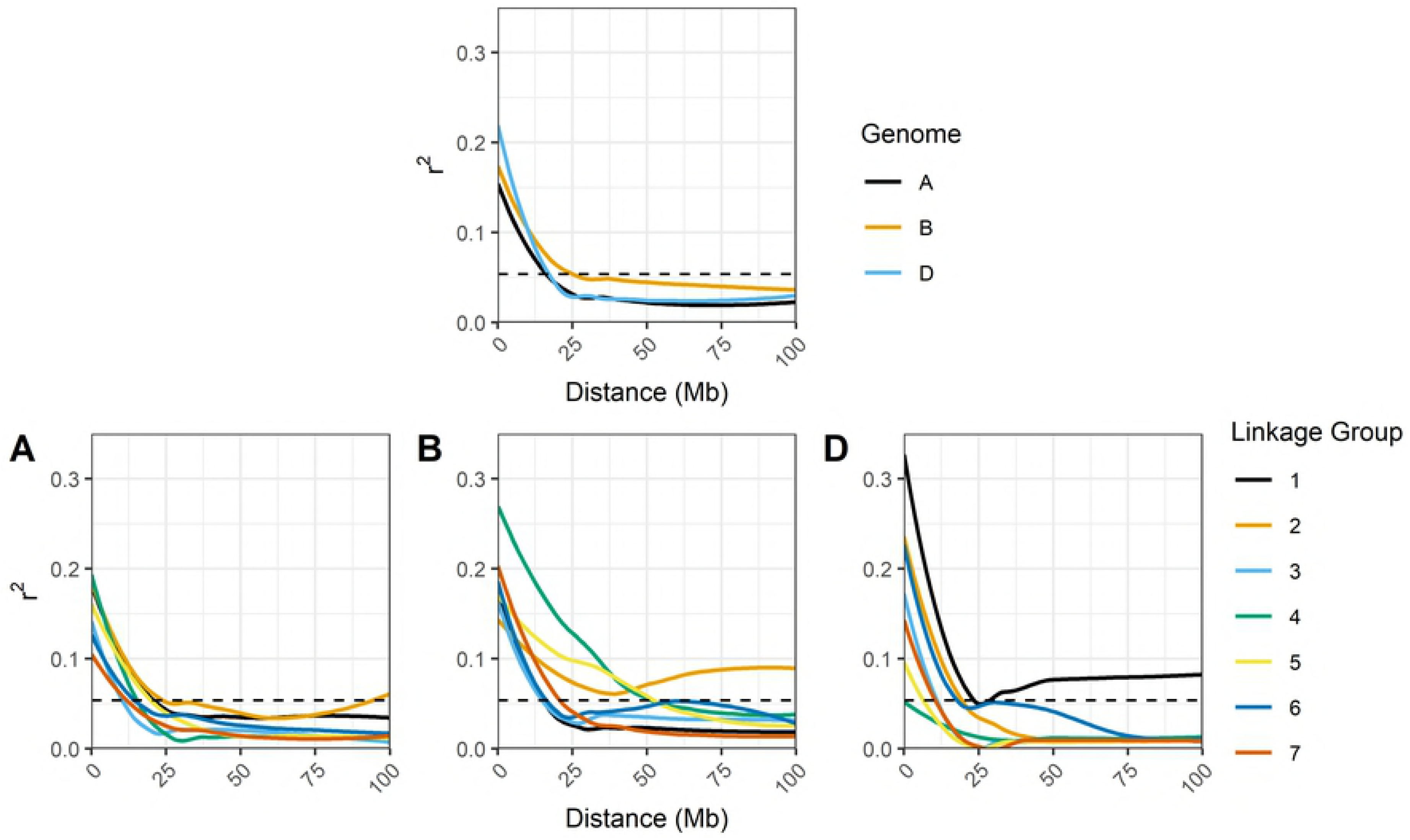
Linkage Disequilibrium by Genome and Chromosome. LOESS regressions of r^2^ between pairs of SNPs vs. physical distance, pooled for each genome (top row), and for each of the seven chromosomes present in the A, B, and D genomes (bottom row).

**Fig 3.**
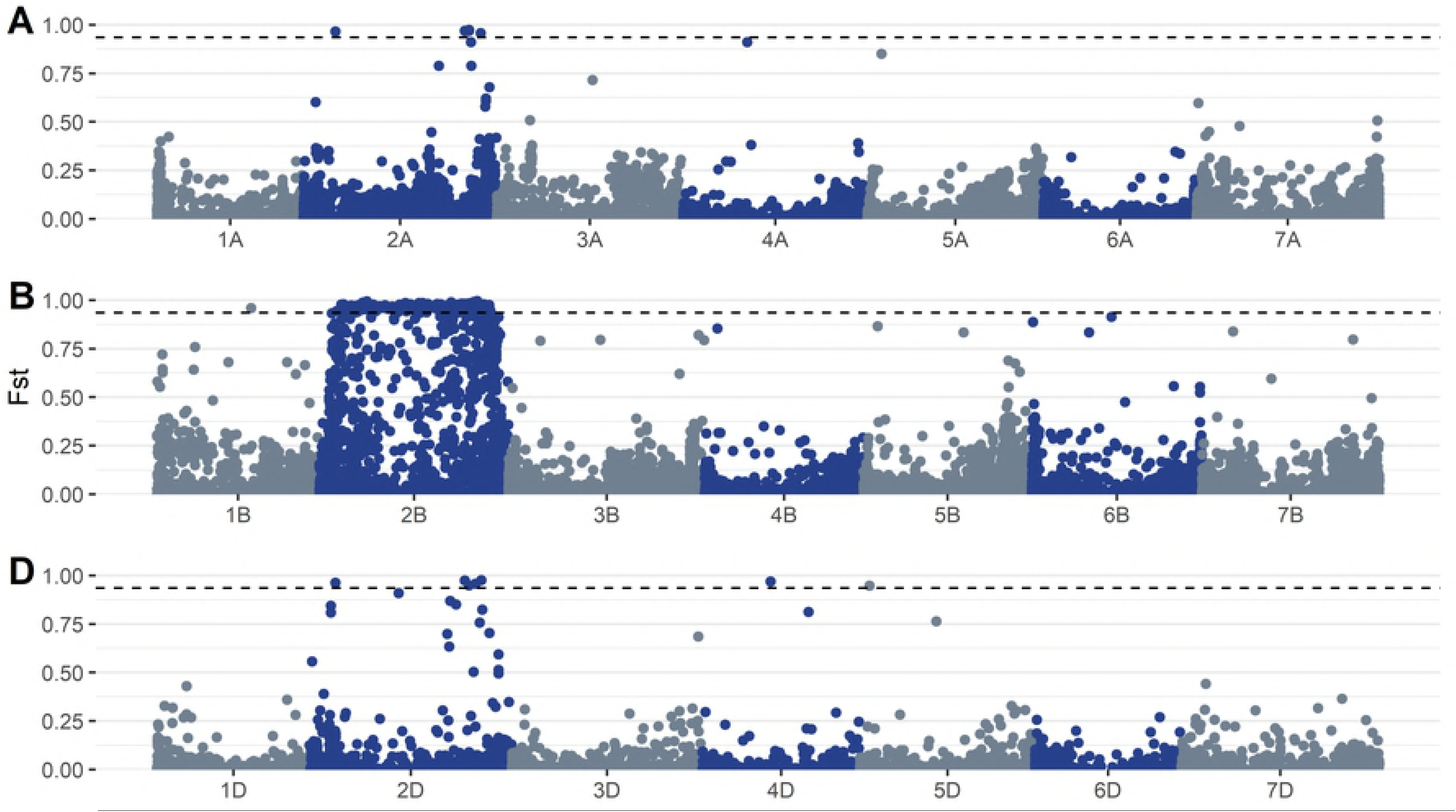
Genome-Wide *F_ST_* Scan. *F_ST_* values were estimated for each SNP in the A genome (top row), B genome (middle row), and D genome (bottom row). The dashed line signifies the 99^th^ percentile of *F_ST_* values.

**Fig 4.**
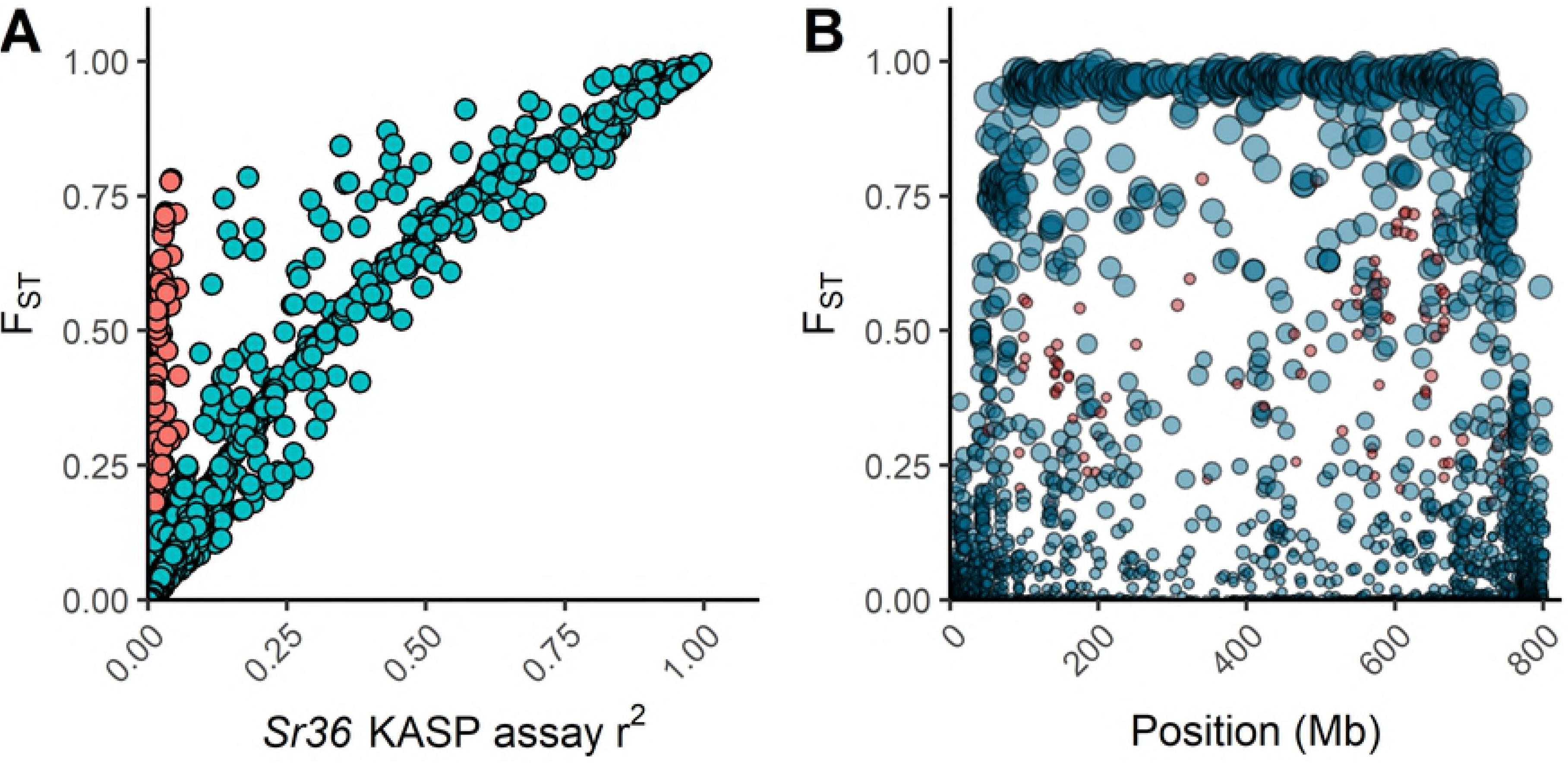
*F_ST_* vs. LD with the *Sr36* KASP Assay. (A) *F_ST_* plotted against r^2^ with the KASP *Sr36* assay for all SNPs on chromosome 2B. Two clusters of SNPs were manually differentiated – those for which *F_ST_* and r^2^ values show a general linear relationship (blue), and those for which *F_ST_* shows no relationship with r^2^ (red). (B) The same clusters of SNPs plotted across chromosome 2B. The size of points indicates the level of r^2^ with the *Sr36* KASP marker.

### Genome-wide association studies

In general, the single-locus model implemented in GCTA identified far fewer significant MTAs than FarmCPU. In total, GCTA identified 44 significant MTAs at the 0.05 FDR significance threshold. However, many of these were located within high-LD blocks and, therefore, clustered at the same putative underlying QTLs. Discounting those SNPs believed to co-localize to identical QTLs yielded nine unique MTAs for HD, MAT, SSQM, TKW, and TWT (**Table 3**). One SNP, represented in the notation *chromosome:position* as 7D:58633321, was pleiotropic for the traits HD and MAT, and hence the nine significant MTAs represented eight putative QTLs. In contrast, FarmCPU identified 108 MTAs at the same significance threshold. Removing those SNPs with RMIPs below 0.1 yielded a total of 74 significant MTAs for the traits FLSG, GSQM, GW, HD, HT, MAT, NDVI, PROT, SPH, SSQM, STARCH, TKW, TWT, and YLD (**Table 4**). After filtering by the RMIP threshold, the remaining significant FarmCPU MTAs exhibited a uniform distribution for RMIP values. A significant MTA affecting the trait TWT located on the long arm of 6A had the highest RMIP value of 0.99.

**Table 3:** Significant marker-trait associations identified by the GCTA leave-one-chromosome-out method.

| Trait<sup>a</sup> | Chr<sup>b</sup> | Pos<sup>b</sup> | cM | Alleles | MAF | P-value | Effect | Units | Coincident Loci<sup>c</sup> |
| --- | --- | --- | --- | --- | --- | --- | --- | --- | --- |
| TKW | 1A | 40498783 | 68 | G\T | 0.09 | 1.62E-06 | 1.25 | grams | W4ZY32 \| Patatin (626Kb) |
| TWT | 1A | 583992305 | 218.8 | T\C | 0.23 | 9.33E-07 | -3.39 | g L<sup>-1</sup> |  |
| HD | 1B | 50989662 | 52.8 | G\A | 0.29 | 1.00E-06 | 0.533 | Julian days | W5A8F0 \| Sugar transporter SWEET (116Kb) |
| MAT | 2B | 3266917 | 0 | C\A | 0.15 | 2.35E-05 | -0.527 | Julian days |  |
| SSQM | 3A | 6788503 | 12.5 | C\T | 0.46 | 1.56E-06 | 16 | Spikes m<sup>-2</sup> | G1UE17 \| *MFT* (506Kb) |
| HD | 6A | 150099119 | 109.5 | A\T | 0.17 | 1.52E-05 | -0.507 | Julian days |  |
| HD | 6A | 542256574 | 128.5 | T\C | 0.34 | 3.39E-05 | -0.417 | Julian days |  |
| HD | 7D | 58633321 | 84.6 | A\G | 0.33 | 3.70E-10 | -0.634 | Julian days |  |
| MAT | 7D | 58633321 | 84.6 | A\G | 0.33 | 5.71E-07 | -0.495 | Julian days |  |

Chr chromosome; Pos physical position; cM genetic position in centiMorgans; Alleles major allele listed first, minor allele second; Effect mean value of lines containing minor allele minus mean value of lines containing major allele; Coincident QTLs loci identified in previous studies believed to co-locate with QTL identified in the present study
<sup>a</sup> HD heading date; MAT physiological maturity date; SSQM spikes per square meter; TKW thousand-kernel weight; TWT test weight
<sup>b</sup> Boxes indicate QTLs with pleiotropic effects
<sup>c</sup> Column formatted as “UniProt Accession ID (if known) | Gene/feature name (Physical Distance or r<sup>2</sup>)”

**Table 4:**
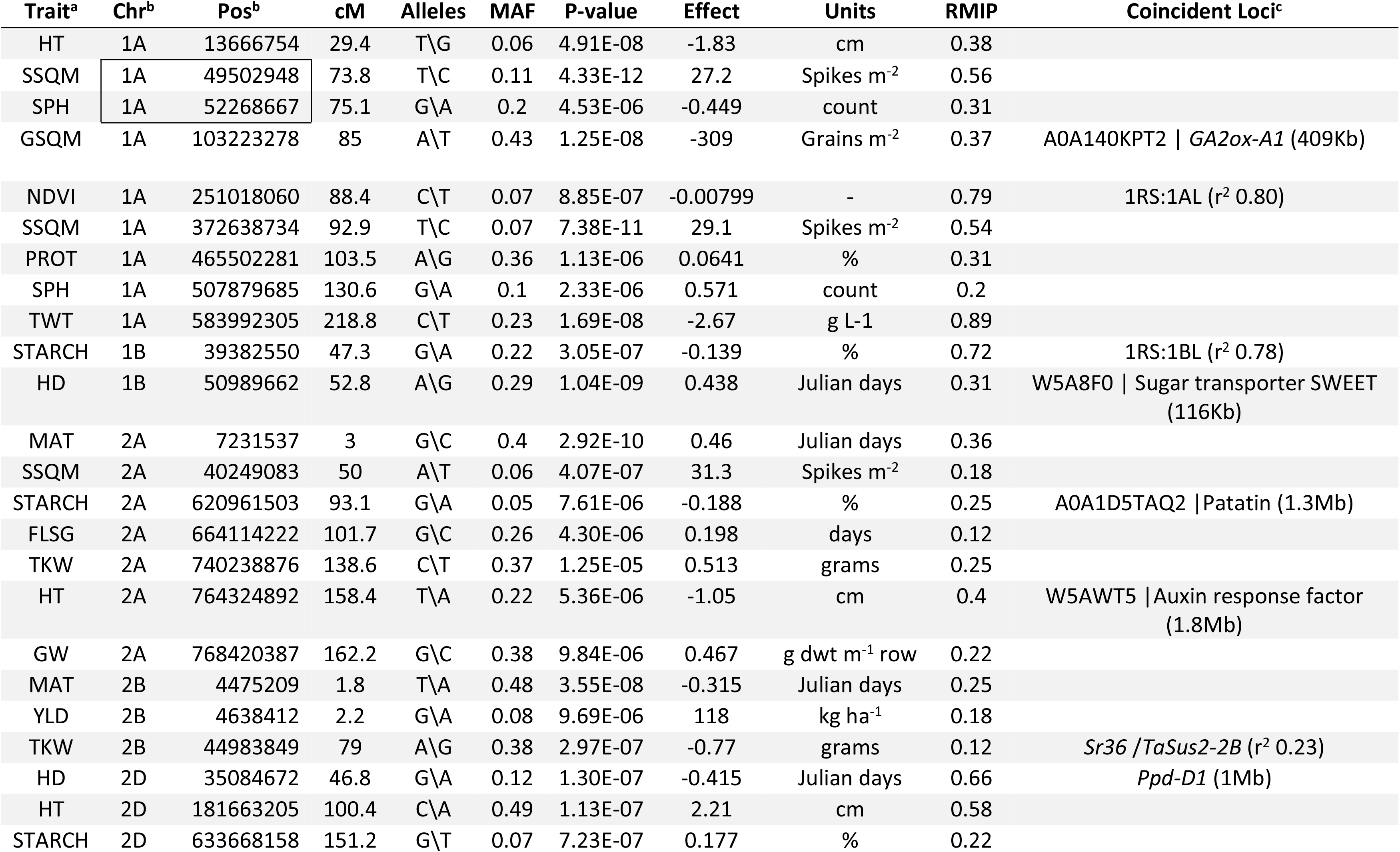
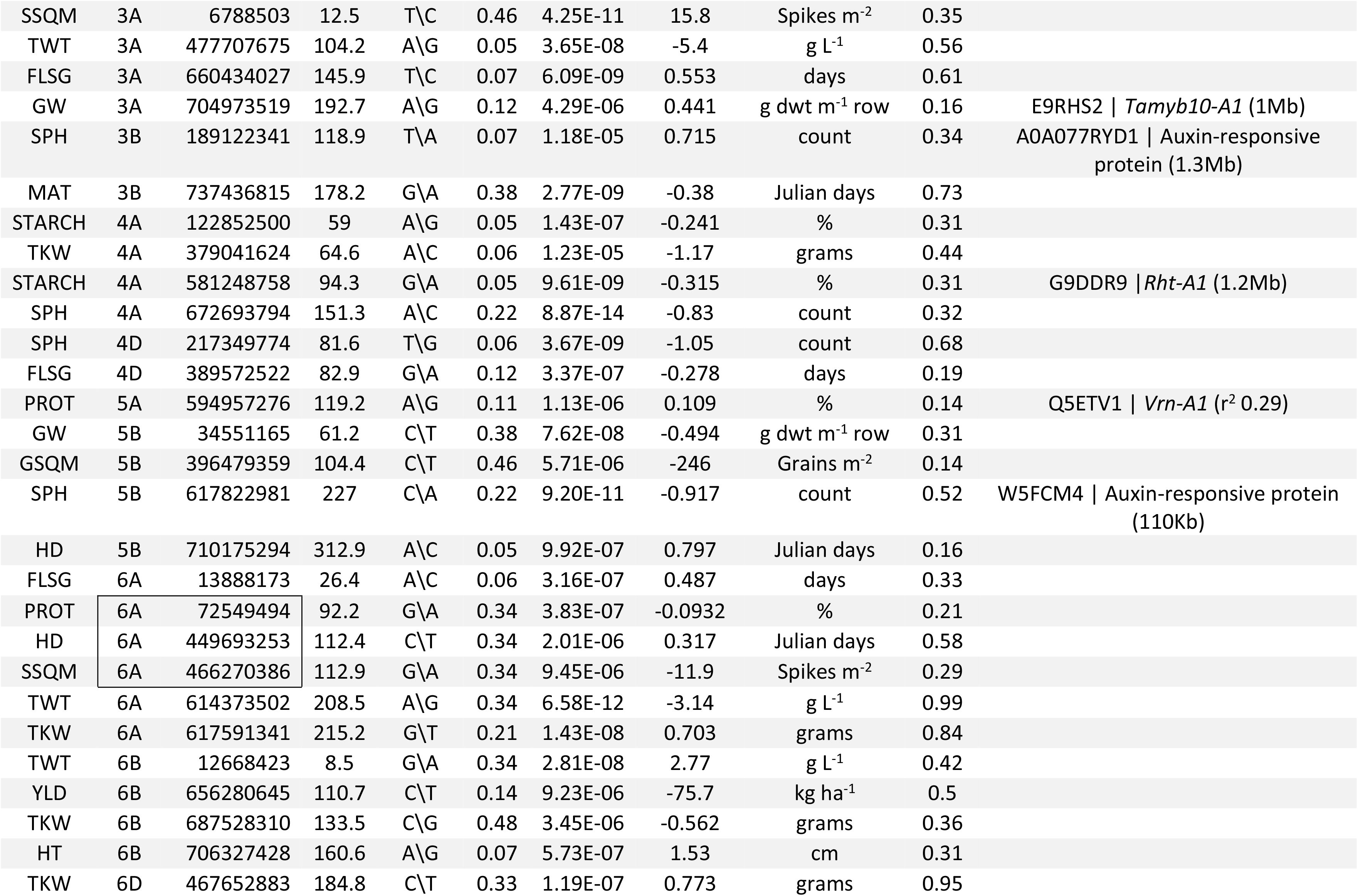
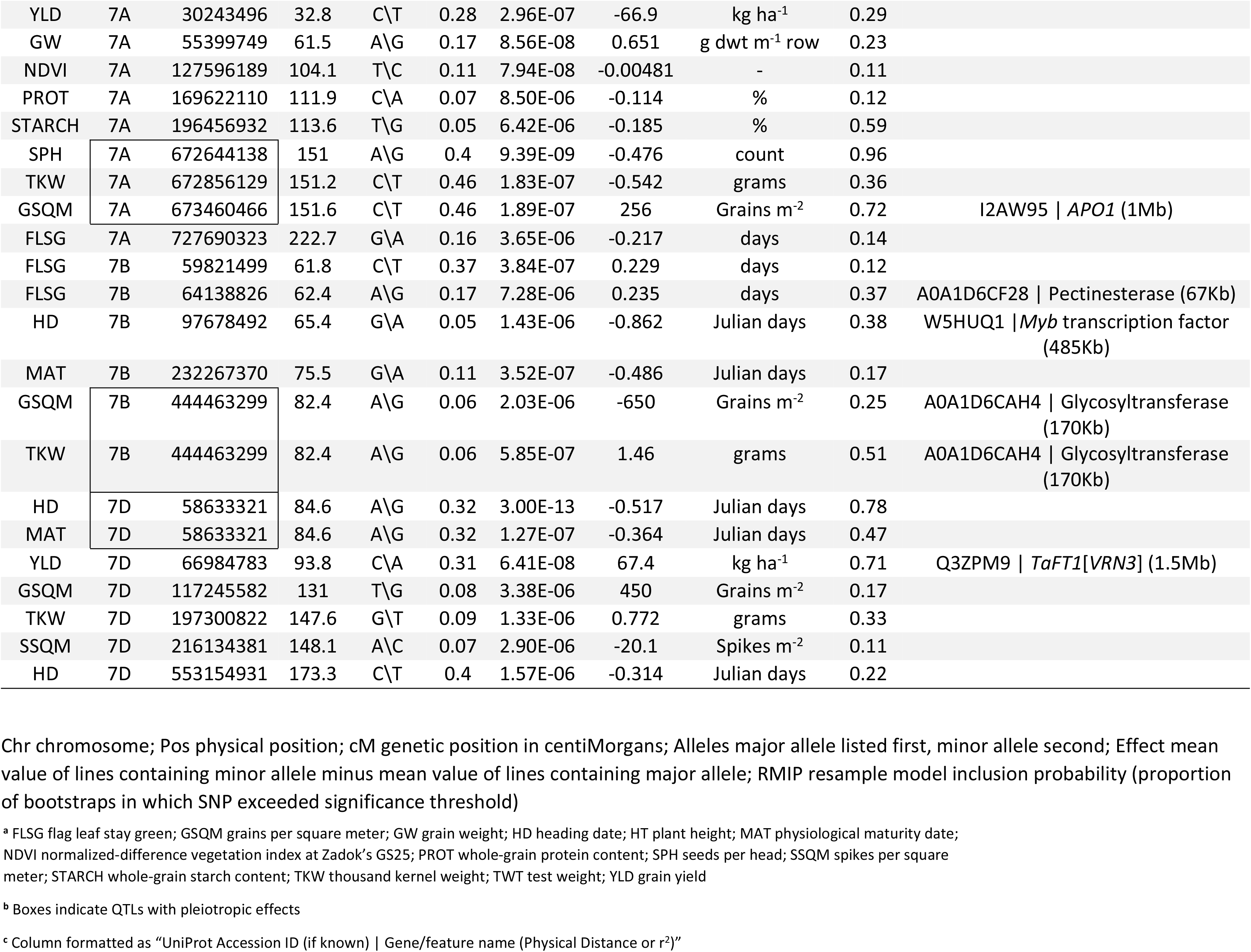
Significant marker-trait associations identified by the FarmCPU algorithm.

Since FarmCPU fit significant MTAs as covariates based partially upon its internal LD estimates, the MTAs identified by this model all reside within different putative QTLs. However, several MTAs did cluster in high-LD regions with pleiotropic effects, and hence the 74 total MTAs represent 67 putative QTLs. Of the eight putative QTLs identified by GCTA, four were also identified by FarmCPU. Examination of the Manhattan plots and uniform quantile-quantile plots of the p-values produced by the single-locus model (**S1 File**) and FarmCPU (**S2 File**) demonstrated adequate control of p-value inflation for most traits, with FarmCPU properly detecting and correcting for LD between SNPs co-located on QTLs.

### MTA haplotypes and translation effects

The GCTA and FarmCPU analyses identified a total of 77 significant MTAs, 35 of which were located on haplotype blocks containing two or more SNPs (**S4 Table-A**). These SNPs overlapped 48 genes encoding 68 transcripts, and produced a total of 109 predicted transcriptional consequences due to alternative splicing (**S4 Table-B**). One haplotype block spanning a 2Mb segment on chromosome 7A contained three significant MTAs (7A: 672644138, 7A: 672856129, and 7A: 673460466). The most severe predicted consequences were isolated and compiled for each SNP. A total of 56 of the SNPs involved in significant MTAs (73%) were intergenic, and of these intergenic SNPs, 15 were labeled as upstream or downstream proximal variants (i.e. within 5Kb of the start or end of a gene). The remaining 21 SNPs (27%) were located within genes, with 11 of these occurring in introns, one occurring in a 5’ untranslated region, two occurring in 3’ untranslated regions, and the remaining seven causing missense mutations within exons. Putative functions for genes containing SNPs causing missense substitutions include a disease resistance protein, a RNA binding protein, a ubiquitin-family protein, a chloroplast membrane translocase, an aspartyl protease family protein, a glutamate receptor, and a *Myb* transcription factor. In addition, a total of 277 wheat genes of known function were located within 2Mb of the set of significant MTAs.

## Discussion

The current study demonstrates that the recent availability of the wheat reference genome will greatly facilitate reliable QTL curation and comparisons between studies. The ability to describe MTAs in terms of both physical and genetic positions removes much of the ambiguity associated with earlier studies which relied solely on genetic marker positions. The availability of a reference genome also enables more accurate QTL and haplotype boundary delineation and comparative genomics analyses. We were able to use positional information both to identify GBS SNPs that could potentially be used as substitutes for existing KASP SNP assays, and to identify candidate genes located near significant MTAs.

In the present study, many of the previously characterized loci of agronomic importance interrogated with KASP-SNP assays had significant effects on multiple traits (**S3 Table-B**). Loci that significantly affected many traits included the *Sr36* stem rust resistance gene and the *TaSus2-2B* sucrose synthase gene (both assaying the presence/absence of the 2G:2B *T. timopheevi* translocation), the 1AL:1RS and 1BL:1RS translocations, the *Ppd-B1* and *Ppd-D1* photosensitivity genes, the *Rht-B1* and *Rht-D1* dwarfing genes, and polymorphisms within the *Vrn-A1* vernalization gene. However, despite their significant effects on many traits, the KASP assays for these loci were not among the significant MTAs identified in the current study. Some potential explanations for these results are presented below.

The two *Rht* dwarfing genes *Rht-B1b* and *Rht-D1b* mostly occurred in repulsion within the testing panel. Of the 324 lines included in the panel, 22 had neither the *Rht-B1b* nor the *Rht-D1b* dwarfing alleles, one line was heterozygous for both alleles, and one line was homozygous for both alleles, with the rest being homozygous for either one dwarfing allele or the other. Note that lines lacking both *Rht-B1b* and *Rht-D1b* may contain other dwarfing genes which were not assayed in this study. At face value, this high degree of repulsion led to the somewhat odd finding that the presence of the *Rht-B1b* allele increased height and decreased yield when allelic effects were calculated for each of the KASP assays individually. A subsequent ANOVA pooling together the two dwarfing alleles revealed that at an alpha level of 0.05 lines with either *Rht-B1b* or *Rht-D1b* were significantly shorter and yielded significantly higher than lines that were wild-type for both genes. Yield was significantly higher for lines with only *Rht-D1b* vs. those with only *Rht-B1b*, though height was not significantly different between lines with these dwarfing genes. (These analyses excluded lines that were heterozygous for either allele, and the single line that was homozygous for both alleles). This perhaps serves as a good reminder of how population-specific inter-locus interactions can prevent the accurate estimation of many allele effects. While multi-locus GWAS models may partially ameliorate these problems in some cases, the careful choice of experimental design, such as the use of nested association mapping panels [63], is likely a better solution.

In the present study, both the *Sr36* and *TaSus2-2B* genes were highly confounded with the testing panel’s population structure. A SNP located at 2B:44983849 was in moderate LD with both *Sr36* and *TaSus2-2B* (r^2^ = 0.24), and had the expected significant impact on TKW, given the known effects that *TaSus2-2B* exerts on kernel weight [13]. The 2G:2B translocation from *T. timopheevi* on which the *Sr36* gene and the *TaSus2-2B-HapG* allele reside introduced a large alien haplotype into the panel germplasm, driving the high LD observed in the B genome and on chromosome 2B specifically (**Fig 2**). The findings regarding LD in the B genome are in contrast to the findings of Chao et al. [64], who reported lower levels of LD in the B genome of both winter and spring wheat. However, it is not known how frequently the 2G:2B translocation occurred within the germplasm they tested. In contrast, Cavanagh et al. [65] found that *Sr36* was associated with a large segment of segregation distortion on chromosome 2B in seven different mapping populations. Splitting the population by the presence or absence of the *Sr36* gene and subsequently performing a genome-wide F_ST_ scan (**Fig 3**) revealed a large high-LD block spanning most of 2B, containing many SNPs which could almost perfectly differentiate between lines with and without the 2G:2B translocation. In addition, an enrichment in high-F_ST_ SNPs was observed on chromosomes 2A and 2D, likely due to SNP misalignments between chromosomes within this homeologous group. On chromosome 2B, there was a general positive linear trend between a SNP’s *F_ST_* score and its LD with the *Sr36* KASP marker. This was as expected, as the presence or absence of this marker was used to partition the subpopulations. However, a subset of SNPs simultaneously exhibited high F_ST_ and an r^2^ value with the KASP *Sr36* assay that approached zero. These SNPs were distributed throughout the translocation (**Fig 4**). It is not immediately apparent what factor is driving this segregation of SNPs, but it is likely that SNPs with a high F_ST_ value that are also in high LD with the *Sr36* KASP marker share a common private haplotype allele inherited from *T. timopheevi*, while those with a high *F_ST_* which are in low LD with the *Sr36* KASP assay represent a set of SNPs with shared alleles between *T. aestivum* and *T. timopheevi*. Despite the effects of the 2G:2B translocation in the current study, population structure was generally subdued, as evidenced by principal component plots of the LD-thinned dataset, where the first principal component explained only 3.68% of total variation (**Fig 1-B**). This finding is in line with those of previous studies examining population structure in elite European winter wheat germplasm [66,67]. This suggests extensive past admixture among the lines included in the population, which is as expected given the frequent germplasm exchanges that are typical of public small grains breeding programs.

A previous report by Sukumaran et al. [21] found much more pronounced population structure effects due to rye translocations, using a panel of elite spring germplasm which clustered into two distinct sub-populations explained by the presence or absence of the 1BL:1RS translocation. In addition, Sukumaran et al. found that the 1BL:1RS translocation explained significant differences among the two subpopulations for most of the traits included in their study (e.g. grain yield, grain number, grain weight, plant height, and several phenological traits). The 1AL:1RS and 1BL:1RS translocations have been associated with desirable disease and insect resistance traits, as well as drought and general environmental stress resistance. However, the 1BL:1RS translocation has been associated with lateness, and effects on yield due to these translocations appear to be more variable, depending upon environment and genetic background [68–70]. While both the 1AL:1RS and 1BL:1RS translocations did produce significant differences for many traits (**S3 Table-B**), their contributions to population structure were not noticeable in comparison to the effects of the 2G:2B translocation (data not shown). In addition, the indicator KASP markers for these translocations did not produce significant p-values in either the GCTA or FarmCPU GWAS. The 1AL:1RS translocation occurred at a very low frequency (0.07) within the tested germplasm, while the 1BL:1RS translocation occurred at a frequency of 0.19. In the study of Sukumaran et al. [21], the 1BL:1RS translocation occurred at a frequency of 0.39. In addition, the KASP markers for 1AL:1RS and 1BL:1RS were in high LD (r^2^ approximately 0.8) with the SNPs 1A:103223278 and 1B:39382550 respectively, which likely masked the effects of the KASP markers. This high LD suggests that these GBS markers, or others in close proximity, could be used as suitable indicators for the 1AL:1RS and 1BL:1RS translocations.

The effects of previously-characterized genes affecting phenological traits remained insignificant in the GWAS due to a variety of factors. Polymorphisms of *Vrn-A1* exon 7 and *Vrn-B1* occurred at low frequencies of 0.03 and 0.07 within the testing panel, respectively, precluding their detection in the GWAS. The *Vrn-A1* exon 4 SNP had a higher minor allele frequency of 0.14. However, this SNP was in moderate LD (r^2^ = 0.29) with one GBS SNP (5A:594957276). In addition, the *Vrn-A1* KASP assays used in this study may simply have not been predictive enough to adequately distinguish among vernalization alleles in the tested germplasm. The *Ppd* loci were likewise never identified as significant, most likely due to masking from the predominance of long-vernalizing genotypes in the panel. However, a BLAST analysis indicated that the GBS SNP 2D:35084672, which FarmCPU identified as significant for the trait HD, was located within approximately 1Mb of *Ppd-D1*.

As previously mentioned, there was evidence for pleiotropic effects involving multiple SNPs and traits in both the GCTA and FarmCPU results (**Tables 3 and 4**). The sole region of pleiotropic effect identified by GCTA was a QTL located at approximately 58.5Mb (84.6 cM) on chromosome 7D, affecting the phenological traits HD and MAT. The haplotype block analysis indicated that this QTL spanned a region of roughly 1.5Mb. This QTL was also identified by FarmCPU, and was one of the more consistently significant QTLs identified, with a RMIP value of 0.78. Many of the other pleiotropic QTLs identified by FarmCPU demonstrated trait compensation effects. For example, the minor allele of a QTL at approximately 50Mb (75cM) on chromosome 1A produced an increase in SSQM, but a slight decrease in SPH. The minor allele of a QTL on chromosome 7A at approximately 673Mb (151cM) caused a decrease in seed size, and an increase in GSQM. It also produced a slight decrease in SPH. The latter finding is against expectations, though the high minor allele frequencies of both of these SNPs (0.4 and 0.46) raise the possibility that the major and minor alleles for one SNP or the other may have switched during processing of the genotypic data. The minor allele of a QTL located at approximately 444Mb (82.4cM) on chromosome 7B conversely produced an increase in TKW and a decrease in GSQM. Finally, a large region on chromosome 6A affected the relatively uncorrelated traits PROT, HD, and SSQM. It is not clear whether this region is being affected by long-range LD, or simply physical linkage. A simple pairwise LD analysis indicated that these SNPs were in moderate LD with each other, though the haplotype block analysis placed all three in separate haplotypes.

Due to the large number of significant findings, we limit further discussion of the GWAS results to several MTAs in close proximity to plausible candidate genes. One cluster of MTAs was situated on the long arm of chromosome 7A, at approximately 673Mb (151cM). This pleiotropic region affected traits related to grain size and grain density per head and unit area (TKW, SPH, and GSQM), spanned a region of approximately 1.5Mb, and overlapped with 22 genes. FarmCPU identified one significant SNP within this block with a RMIP value of 0.96, and another SNP producing a predicted missense protein translation effect in a predicted ortholog of an aspartyl protease family protein. Notably, the haplotype block containing this SNP also contains an ortholog of the aberrant panicle organization 1 (*APO1*) protein, which plays an important role in panicle architecture and development in rice (*Oryza sativa* L.). The loss of function of *APO1* results in rice plants which produce small panicles with a lower number of inflorescence branches and spikelets [71].

Chromosome 7B contained a large number of putative candidate genes for multiple traits. A pectin esterase was located in close proximity (67Kb) with a MTA affecting FLSG located at 64Mb (62.4cM). Due to the ubiquity of pectin within plant cell walls, pectin esterases have been associated with a wide range of cellular processes (reviewed in [72]). Notably, pectin esterases have been found to play a role in fruit maturation in heat-stressed tomato plants [73], as well as the initiation of flowering in day lilies [74]. Another MTA on 7B located at 97Mb (65.4cM) was within 485Kb of a MYB transcription factor. The MYB family of proteins function in a wide number of signal transduction pathways in plants. Gibberellin-interacting MYB factors (GAMYBs) were first identified in barley (*Hordeum vulgare* L.), inducing the activation of alpha-amylase in the aleurone tissue of grains [75]. GAMYB factors were subsequently identified in wheat [76]. Currently, GAMYB factors have been putatively implicated in contributing to flowering initiation, though their role in this process is not well understood. Gocal et al. [77] suggested that a GAMYB protein in the grass species *Lolium temulentum* L. was an important component in signaling pathways for flower development. However, Kaneko et al. [78] later found that while rice GAMYB was important in normal flower organ development, knockout mutants did not display any differences from wild-type plants in the timing of flower development. The pleiotropic SNP located at approximately 444Mb (82.4cM) on chromosome 7B exerted large effects on the traits GSQM and TKW, and was intronic within an ortholog of an intracellular protein transporter identified in *Medicago truncatula* Gaertn. While this SNP is not associated with a haplotype block, it is located 170Kb from a glycosyltransferase gene. The glycosyltransferases are a superfamily consisting of thousands of identified proteins. Notably, multiple families of glycosyltransferases have been associated with grain development in wheat [79].

Finally, a SNP located at approximately 59Mb (84.6cM) on chromosome 7D was identified by both the single-locus and FarmCPU models, produced pleiotropic effects for both HD and MAT, had a large effect size magnitude of approximately 0.5 days, and produced a RMIP value of 0.78 for HD. Due to its exertion of effects with the same sign on both HD and MAT, it is likely that this locus could be involved with plant vernalization or photoperiod response. While this SNP is approximately 10Mb away from the *VRN3* vernalization response/flowering time gene, the significant MTA 7D:66984783 affecting yield is located only 1.5Mb away from *VRN3*, and there is no significant LD between these two SNPs, making it likely that some other as yet undetermined causal agent is underlying this locus’ effects on heading and maturation dates. This SNP is intronic within an ortholog of the *Escherichia coli* Acetyl-coenzyme A carboxylase carboxyl transferase subunit alpha gene, which is involved in fatty acid synthesis via the synthesis of malonyl-CoA [80], though any connection between this function and vernalization response in plants remains unknown.

Common gene ontologies for wheat genes overlapping or proximal to significant MTAs included auxin-interacting proteins and auxin-response transcription factors. These proteins were in close proximity to several MTAs for the trait SPH, located on chromosomes 3B, 5B, and the previously-mentioned pleiotropic region on 7A. One auxin response transcription factor also occurred near a MTA affecting HT on chromosome 2A. Proteins containing a patatin-like phospholipase domain were located near a MTA affecting TKW on chromosome 1A, identified by the single-locus model, and a MTA affecting STARCH on chromosome 2A, identified by FarmCPU. Patatin is the primary storage glycoprotein in potato (*Solanum tuberosum* L.) tubers [81]. While the connection between these patatin-like proteins and the traits TKW and STARCH are speculative, the *dense and erect panicle 3* (*DEP3*) protein in rice contains a patatin-like phospholipase domain, and has been associated with higher-yielding cultivars [82]. Erect panicle rice cultivars exhibit higher yields due to a variety of factors, including panicle architecture, higher lodging resistance, accelerated carbon dioxide diffusion, and better light interception [83].

While the availability of a reference genome allows for previously impossible analyses, this study also identifies a number of areas for potential improvement for GWAS analyses in wheat. For instance, the genome-wide *F_ST_* scan based upon the presence or absence of *Sr36* revealed multiple SNPs which were misaligned between the group 2 homeologous chromosomes, and therefore non-allelic. In addition to the obvious problem of potentially identifying significant MTAs on the wrong chromosome, these SNPs complicate the process of haplotype estimation, as they may artificially “break” patterns of strong LD within haplotype blocks. There is currently no simple solution to identify misaligned SNPs within GBS datasets, though increasingly reliable methods for identifying non-allelic SNPs in polyploids are under development (e.g. [84,85]). Finally, although the availability of a reference genome has greatly aided the interpretation of GWAS studies in wheat, it should be noted that the causal variants underlying the majority of significant MTAs identified in this study remain unknown. As the amount of bioinformatics data available for wheat increases, this situation may improve through the use of new techniques such as gene set analyses.

## Conclusions

The significant MTAs reported in this study indicate that there is still genetic variation in the tested elite germplasm that may be exploited for yield gains. In particular, the combination of identified MTAs affecting traits relating to grains per unit area and phenological development offer promise for increasing the former while avoiding the penalizing effect of lower average grain weights, as suggested in previous literature. In addition, this study suggests that GBS markers can be used to capture much of the variance explained by previously-characterized polymorphisms of major effect. We made use of the first reference genome assembled for wheat, enabling the identification of MTAs based on both physical and genetic positions; it is hoped that the ability to anchor MTAs by physical position will lead to better curation of results and consistency across GWAS studies in the future. This study also identifies some potential targets for future *in vitro* studies to ascertain the biological functions of several candidate genes affecting yield-related traits in wheat. Future challenges will include the proper design of GBS or other genotyping assays to capture the effects of previously-characterized polymorphisms while simultaneously allowing for the discovery of novel polymorphisms affecting traits of interest, better identification of non-allelic SNPs which are misaligned between homeologous chromosomes, and more thorough characterization of gene functions to ease the identification of candidate genes following association analyses.

## Supporting Information

**S1 Table**. List of germplasm tested in the study.

**S2 Table**. Description of traits examined in the study, with trait ontologies as described in http://www.planteome.org/

**S3 Table**. Design of KASP SNP assays for interrogating previously-characterized loci of major effect, and a summary of the allelic effects of these loci.

**S4 Table**. List of genes overlapping significant SNPs and haplotype blocks, predicted protein translation effects for all significant SNPs, and list of all wheat genes with annotations in UniProt occurring within 2Mb of significant MTAs.

**S1 Fig**. Individual and cumulative portions of variance explained by the first 25 principal components of the imputed genotypic data, prior to LD-based filtering.

**S1 File**. Manhattan and QQ plots for SNP p-values generated by the GCTA leave-one-chromosome-out (LOCO) mixed linear model GWAS

**S2 File**. Manhattan and QQ plots for SNP p-values generated by the FarmCPU GWAS

## References

1. Food and Agriculture Organization of the UN. FAOSTAT Food and Agricultural Commodities Production [Internet]. 2013 [cited 30 Mar 2016]. Available: http://faostat3.fao.org/

2. USDA Foreign Agricultural Service. USDA Production, Supply and Distribution Online [Internet]. 2015 [cited 30 Mar 2016]. Available: https://apps.fas.usda.gov/psdonline/

3. Bruinsma J. The resources outlook: by how much do land, water and crop yields need to increase by 2050? Looking Ahead in World Food and Agriculture: Perspectives to 2050. Rome, Italy: Food and Agriculture Organization of the United Nations; 2011. pp. 233–278.

4. Sharma RC, Crossa J, Velu G, Huerta-Espino J, Vargas M, Payne TS, et al. Genetic gains for grain yield in CIMMYT spring bread wheat across international environments. Crop Sci. 2012;52: 1522–1533. doi:10.2135/cropsci2011.12.0634

5. Green AJ, Berger G, Griffey CA, Pitman R, Thomason W, Balota M, et al. Genetic yield improvement in soft red winter wheat in the eastern United States from 1919 to 2009. Crop Sci. 2012;52: 2097–2108. doi:10.2135/cropsci2012.01.0026

6. Donald CM. The breeding of crop ideotypes. Euphytica. 1968;17: 385–403. doi:10.1007/BF00056241

7. Slafer GA. Genetic basis of yield as viewed from a crop physiologist’s perspective. Ann Appl Biol. 2003;142: 117–128. doi:10.1111/j.1744-7348.2003.tb00237.x

8. Borrás L, Slafer GA, Otegui ME. Seed dry weight response to source-sink manipulations in wheat, maize and soybean: a quantitative reappraisal. Field Crops Res. 2004;86: 131–146. doi:10.1016/j.fcr.2003.08.002

9. Reynolds MP, Pellegrineschi A, Skovmand B. Sink-limitation to yield and biomass: a summary of some investigations in spring wheat. Ann Appl Biol. 2005;146: 39–49. doi:10.1111/j.1744-7348.2005.03100.x

10. Slafer G, Calderini D, Miralles D. Yield components and compensation in wheat: opportunities for further increasing yield potential. Increasing yield potential in wheat: Breaking the Barriers. 1996. pp. 101–133.

11. Silva SA, Carvalho FIF de, Caetano V da R, Oliveira AC de, Coimbra JLM de, Vasconcellos NJS de, et al. Genetic basis of stay-green trait in bread wheat. J New Seeds. 2001;2: 55–68. doi:10.1300/J153v02n02_05

12. Guedira M, Xiong M, Hao YF, Johnson J, Harrison S, Marshall D, et al. Heading date QTL in winter wheat (*Triticum aestivum* L.) coincide with major developmental genes *VERNALIZATION1* and *PHOTOPERIOD1*. PLOS ONE. 2016;11: e0154242. doi:10.1371/journal.pone.0154242

13. Jiang Q, Hou J, Hao C, Wang L, Ge H, Dong Y, et al. The wheat (*T. aestivum*) sucrose synthase 2 gene (*TaSus2*) active in endosperm development is associated with yield traits. Funct Integr Genomics. 2011;11: 49–61. doi:10.1007/s10142-010-0188-x

14. Su Z, Hao C, Wang L, Dong Y, Zhang X. Identification and development of a functional marker of *TaGW2* associated with grain weight in bread wheat (*Triticum aestivum* L.). Theor Appl Genet. 2011;122: 211–223. doi:10.1007/s00122-010-1437-z

15. Ma D, Yan J, He Z, Wu L, Xia X. Characterization of a cell wall invertase gene *TaCwi-A1* on common wheat chromosome 2A and development of functional markers. Mol Breed. 2012;29: 43–52. doi:10.1007/s11032-010-9524-z

16. Myles S, Peiffer J, Brown PJ, Ersoz ES, Zhang Z, Costich DE, et al. Association mapping: critical considerations shift from genotyping to experimental design. Plant Cell Online. 2009;21: 2194–2202. doi:10.1105/tpc.109.068437

17. Breseghello F, Sorrells ME. Association mapping of kernel size and milling quality in wheat (*Triticum aestivum* L.) cultivars. Genetics. 2006;172: 1165–1177. doi:10.1534/genetics.105.044586

18. Neumann K, Kobiljski B, Denčić S, Varshney RK, Börner A. Genome-wide association mapping: a case study in bread wheat (*Triticum aestivum* L.). Mol Breed. 2011;27: 37–58. doi:10.1007/s11032-010-9411-7

19. Dodig D, Zoric M, Kobiljski B, Savic J, Kandic V, Quarrie S, et al. Genetic and association mapping study of wheat agronomic traits under contrasting water regimes. Int J Mol Sci. 2012;13: 6167–6188. doi:10.3390/ijms13056167

20. Tadesse W, Ogbonnaya FC, Jighly A, Sanchez-Garcia M, Sohail Q, Rajaram S, et al. Genome-wide association mapping of yield and grain quality traits in winter wheat genotypes. PLoS ONE. 2015;10: e0141339. doi:10.1371/journal.pone.0141339

21. Sukumaran S, Dreisigacker S, Lopes M, Chavez P, Reynolds MP. Genome-wide association study for grain yield and related traits in an elite spring wheat population grown in temperate irrigated environments. Theor Appl Genet. 2015;128: 353–363. doi:10.1007/s00122-014-2435-3

22. Spindel J, Begum H, Akdemir D, Virk P, Collard B, Redoña E, et al. Genomic selection and association mapping in rice (*Oryza sativa*): effect of trait genetic architecture, training population composition, marker number and statistical model on accuracy of rice genomic selection in elite, tropical rice breeding lines. PLoS Genet. 2015;11: e1004982. doi:10.1371/journal.pgen.1004982

23. Salvi S, Tuberosa R. The crop QTLome comes of age. Curr Opin Biotechnol. 2015;32: 179–185. doi:10.1016/j.copbio.2015.01.001

24. The International Wheat Genome Sequencing Consortium (IWGSC), IWGSC RefSeq principal investigators:, Appels R, Eversole K, Feuillet C, Keller B, Rogers J, et al. Shifting the limits in wheat research and breeding using a fully annotated reference genome. Science. 2018;361: eaar7191. doi:10.1126/science.aar7191

25. Zadoks JC, Chang TT, Konzak CF. A decimal code for the growth stages of cereals. Weed Res. 1974;14: 415–421. doi:10.1111/j.1365-3180.1974.tb01084.x

26. Muñoz-Huerta RF, Guevara-Gonzalez RG, Contreras-Medina LM, Torres-Pacheco I, Prado-Olivarez J, Ocampo-Velazquez RV. A review of methods for sensing the nitrogen status in plants: advantages, disadvantages and recent advances. Sensors. 2013;13: 10823–10843. doi:10.3390/s130810823

27. Alley MM, Brann DE, Stromberg EL, Hagood ES, Herbert A, Jones EC, et al. Intensive soft red winter wheat production: a management guide. Blacksburg, VA: Virginia Tech Cooperative Extension Distribution Center; 1993.

28. Phillips SB, Keahey DA, Warren JG, Mullins GL. Estimating winter wheat tiller density using spectral reflectance sensors for early-spring, variable-rate nitrogen applications. Agron J. 2004;96: 591. doi:10.2134/agronj2004.0591

29. AOAC International. Protein (crude) in animal feed (990.03). Official Methods of Analysis, 17th Edition. 17th ed. Rockville, MD: Association of Official Analytical Chemists; 2000.

30. Eilers PHC, Marx BD. Multivariate calibration with temperature interaction using twodimensional penalized signal regression. Chemom Intell Lab Syst. 2003;66: 159–174. doi:10.1016/S0169-7439(03)00029-7

31. R Core Team. R: A Language and Environment for Statistical Computing. [Internet]. R Foundation for Statistical Computing, Vienna, Austria; 2015. Available: http://www.R-project.org/

32. Rodríguez-Álvarez MX, Boer MP, van Eeuwijk FA, Eilers PHC. Correcting for spatial heterogeneity in plant breeding experiments with P-splines. Spat Stat. 2018;23: 52–71. doi:10.1016/j.spasta.2017.10.003

33. Cullis BR, Smith AB, Coombes NE. On the design of early generation variety trials with correlated data. J Agric Biol Environ Stat. 2006;11: 381. doi:10.1198/108571106×154443

34. Bates D, Mächler M, Bolker B, Walker S. Fitting linear mixed-effects models using lme4. J Stat Softw. 2015;67. doi:10.18637/jss.v067.i01

35. Poland JA, Brown PJ, Sorrells ME, Jannink J-L. Development of high-density genetic maps for barley and wheat using a novel two-enzyme genotyping-by-sequencing approach. PLoS ONE. 2012;7: e32253. doi:10.1371/journal.pone.0032253

36. Bradbury PJ, Zhang Z, Kroon DE, Casstevens TM, Ramdoss Y, Buckler ES. TASSEL: Software for association mapping of complex traits in diverse samples. Bioinformatics. 2007;23: 2633–2635. doi:10.1093/bioinformatics/btm308

37. Glaubitz JC, Casstevens TM, Lu F, Harriman J, Elshire RJ, Sun Q, et al. TASSEL-GBS: A high capacity genotyping by sequencing analysis pipeline. PLOS ONE. 2014;9: e90346. doi:10.1371/journal.pone.0090346

38. Li H, Durbin R. Fast and accurate short read alignment with Burrows–Wheeler transform. Bioinformatics. 2009;25: 1754–1760. doi:10.1093/bioinformatics/btp324

39. Browning SR, Browning BL. Rapid and accurate haplotype phasing and missing-data inference for whole-genome association studies by use of localized haplotype clustering. Am J Hum Genet. 2007;81: 1084–1097. doi:10.1086/521987

40. Chang CC, Chow CC, Tellier LC, Vattikuti S, Purcell SM, Lee JJ. Second-generation PLINK: rising to the challenge of larger and richer datasets. GigaScience. 2015;4: 7. doi:10.1186/s13742-015-0047-8

41. Chen Y, Carver BF, Wang S, Zhang F, Yan L. Genetic loci associated with stem elongation and winter dormancy release in wheat. Theor Appl Genet. 2009;118: 881–889. doi:10.1007/s00122-008-0946-5

42. Díaz A, Zikhali M, Turner AS, Isaac P, Laurie DA. Copy number variation affecting the *Photoperiod-B1* and *Vernalization-A1* genes is associated with altered flowering time in wheat (*Triticum Aestivum*). PLOS ONE. 2012;7: e33234. doi:10.1371/journal.pone.0033234

43. Li G, Yu M, Fang T, Cao S, Carver BF, Yan L. Vernalization requirement duration in winter wheat is controlled by *Ta*VRN-A1 at the protein level. Plant J. 2013;76: 742–753. doi:10.1111/tpj.12326

44. Kippes N, Debernardi JM, Vasquez-Gross HA, Akpinar BA, Budak H, Kato K, et al. Identification of the *VERNALIZATION 4* gene reveals the origin of spring growth habit in ancient wheats from South Asia. Proc Natl Acad Sci. 2015;112: E5401–E5410. doi:10.1073/pnas.1514883112

45. Brown-Guedira GL, Badaeva ED, Gill BS, Cox TS. Chromosome substitutions of *Triticum timopheevii* in common wheat and some observations on the evolution of polyploid wheat species. Theor Appl Genet. 1996;93: 1291–1298. doi:10.1007/BF00223462

46. Nyquist WE. Differential fertilization in the inheritance of stem rust resistance in hybrids involving a common wheat strain derived from *Triticum timopheevi*. Genetics. 1962;47: 1109–1124.

47. Cabrera A, Guttieri M, Smith N, Souza E, Sturbaum A, Hua D, et al. Identification of milling and baking quality QTL in multiple soft wheat mapping populations. Theor Appl Genet. 2015;128: 2227–2242. doi:10.1007/s00122-015-2580-3

48. Zheng X, Levine D, Shen J, Gogarten SM, Laurie C, Weir BS. A high-performance computing toolset for relatedness and principal component analysis of SNP data. Bioinformatics. 2012;28: 3326–3328. doi:10.1093/bioinformatics/bts606

49. Cleveland WS. Robust locally weighted regression and smoothing scatterplots. J Am Stat Assoc. 1979;74: 829–836. doi:10.1080/01621459.1979.10481038

50. Weir BS, Cockerham CC. Estimating F-statistics for the analysis of population structure. Evolution. 1984;38: 1358–1370. doi:10.2307/2408641

51. Danecek P, Auton A, Abecasis G, Albers CA, Banks E, DePristo MA, et al. The variant call format and VCFtools. Bioinformatics. 2011;27: 2156–2158. doi:10.1093/bioinformatics/btr330

52. Yang J, Lee SH, Goddard ME, Visscher PM. GCTA: a tool for genome-wide complex trait analysis. Am J Hum Genet. 2011;88: 76–82. doi:10.1016/j.ajhg.2010.11.011

53. Benjamini Y, Hochberg Y. Controlling the false discovery rate: a practical and powerful approach to multiple testing. J R Stat Soc Ser B Methodol. 1995;57: 289–300.

54. Liu X, Huang M, Fan B, Buckler ES, Zhang Z. Iterative usage of fixed and random effect models for powerful and efficient genome-wide association studies. PLOS Genet. 2016;12: e1005767. doi:10.1371/journal.pgen.1005767

55. Kusmec A, Schnable PS. FarmCPUpp: Efficient large-scale genomewide association studies. Plant Direct. 2018;2: e00053. doi:10.1002/pld3.53

56. Wallace JG, Zhang X, Beyene Y, Semagn K, Olsen M, Prasanna BM, et al. Genome-wide association for plant height and flowering time across 15 tropical maize populations under managed drought stress and well-watered conditions in sub-Saharan Africa. Crop Sci. 2016;56: 2365–2378. doi:10.2135/cropsci2015.10.0632

57. Valdar W, Holmes CC, Mott R, Flint J. Mapping in structured populations by resample model averaging. Genetics. 2009;182: 1263–1277. doi:10.1534/genetics.109.100727

58. Gabriel SB, Schaffner SF, Nguyen H, Moore JM, Roy J, Blumenstiel B, et al. The structure of haplotype blocks in the human genome. Science. 2002;296: 2225–2229. doi:10.1126/science.1069424

59. McLaren W, Pritchard B, Rios D, Chen Y, Flicek P, Cunningham F. Deriving the consequences of genomic variants with the Ensembl API and SNP Effect Predictor. Bioinforma Oxf Engl. 2010;26: 2069–2070. doi:10.1093/bioinformatics/btq330

60. Groos C, Robert N, Bervas E, Charmet G. Genetic analysis of grain protein-content, grain yield and thousand-kernel weight in bread wheat. Theor Appl Genet. 2003;106: 1032–1040. doi:10.1007/s00122-002-1111-1

61. Cox MC, Qualset CO, Rains DW. Genetic variation for nitrogen assimilation and translocation in wheat. I. Dry matter and nitrogen accumulation. Crop Sci. 1985;25: 430–435. doi:10.2135/cropsci1985.0011183X002500030002x

62. Terman GL, Ramig RE, Dreier AF, Olson RA. Yield-protein relationships in wheat grain, as affected by nitrogen and water. Agron J. 1969;61: 755–759. doi:10.2134/agronj1969.00021962006100050031x

63. Yu J, Holland JB, McMullen MD, Buckler ES. Genetic design and statistical power of nested association mapping in maize. Genetics. 2008;178: 539–551. doi:10.1534/genetics.107.074245

64. Chao S, Dubcovsky J, Dvorak J, Luo M-C, Baenziger SP, Matnyazov R, et al. Population-and genome-specific patterns of linkage disequilibrium and SNP variation in spring and winter wheat (*Triticum aestivum* L.). BMC Genomics. 2010;11: 727. doi:10.1186/1471-2164-11-727

65. Cavanagh CR, Chao S, Wang S, Huang BE, Stephen S, Kiani S, et al. Genome-wide comparative diversity uncovers multiple targets of selection for improvement in hexaploid wheat landraces and cultivars. Proc Natl Acad Sci U S A. 2013;110: 8057–8062. doi:10.1073/pnas.1217133110

66. Reif JC, Gowda M, Maurer HP, Longin CFH, Korzun V, Ebmeyer E, et al. Association mapping for quality traits in soft winter wheat. Theor Appl Genet. 2011;122: 961–970. doi:10.1007/s00122-010-1502-7

67. Würschum T, Langer SM, Longin CFH, Korzun V, Akhunov E, Ebmeyer E, et al. Population structure, genetic diversity and linkage disequilibrium in elite winter wheat assessed with SNP and SSR markers. Theor Appl Genet. 2013;126: 1477–1486. doi:10.1007/s00122-013-2065-1

68. Moreno-Sevilla B, Baenziger PS, Peterson CJ, Graybosch RA, McVey DV. The 1BL/1RS translocation: agronomic performance of F3-derived lines from a winter wheat cross. Crop Sci. 1995;35: 1051–1055. doi:10.2135/cropsci1995.0011183X003500040022x

69. Singh RP, Huerta-Espino J, Rajaram S, Crossa J. Agronomic effects from chromosome translocations 7DL.7AG and 1BL.1RS in spring wheat. Crop Sci. 1998;38: 27–33. doi:10.2135/cropsci1998.0011183X003800010005x

70. Ehdaie B, Whitkus RW, Waines JG. Root biomass, water-use efficiency, and performance of wheat–rye translocations of chromosomes 1 and 2 in spring bread wheat ‘Pavon.’ Crop Sci. 2003;43: 710–717. doi:10.2135/cropsci2003.7100

71. Ikeda K, Ito M, Nagasawa N, Kyozuka J, Nagato Y. Rice *ABERRANT PANICLE ORGANIZATION 1*, encoding an F-box protein, regulates meristem fate. Plant J. 2007;51: 1030–1040. doi:10.1111/j.1365-313X.2007.03200.x

72. Micheli F. Pectin methylesterases: Cell wall enzymes with important roles in plant physiology. Trends Plant Sci. 2001;6: 414–419. doi:10.1016/S1360-1385(01)02045-3

73. Kagan-Zur V, Tieman DM, Marlow SJ, Handa AK. Differential regulation of polygalacturonase and pectin methylesterase gene expression during and after heat stress in ripening tomato (*Lycopersicon esculentum* Mill.) fruits. Plant Mol Biol. 1995;29: 1101–1110. doi:10.1007/BF00020455

74. Panavas T, Reid PD, Rubinstein B. Programmed cell death of daylily petals: activities of wall-based enzymes and effects of heat shock. Plant Physiol Biochem. 1998;36: 379–388. doi:10.1016/S0981-9428(98)80079-X

75. Gubler F, Kalla R, Roberts JK, Jacobsen JV. Gibberellin-regulated expression of a *myb* gene in barley aleurone cells: evidence for Myb transactivation of a high-pI α-amylase gene promoter. Plant Cell. 1995;7: 1879–1891. doi:10.1105/tpc.7.11.1879

76. Chen L, Nishizawa T, Higashitani A, Suge H, Wakui Y, Takeda K, et al. A variety of wheat tolerant to deep-seeding conditions: elongation of the first internode depends on the response to gibberellin and potassium. Plant Cell Environ. 2001;24: 469–476. doi:10.1046/j.1365-3040.2001.00688.x

77. Gocal GFW, Poole AT, Gubler F, Watts RJ, Blundell C, King RW. Long-day up-regulation of a *GAMYB* gene during *Lolium temulentum* inflorescence formation. Plant Physiol. 1999;119: 1271–1278. doi:10.1104/pp.119.4.1271

78. Kaneko M, Inukai Y, Ueguchi-Tanaka M, Itoh H, Izawa T, Kobayashi Y, et al. Loss-of-function mutations of the rice *GAMYB* gene impair α-amylase expression in aleurone and flower development. Plant Cell. 2004;16: 33–44. doi:10.1105/tpc.017327

79. Sado P-E, Tessier D, Vasseur M, Elmorjani K, Guillon F, Saulnier L. Integrating genes and phenotype: a wheat–*Arabidopsis*–rice glycosyltransferase database for candidate gene analyses. Funct Integr Genomics. 2009;9: 43–58. doi:10.1007/s10142-008-0100-0

80. Freiberg C, Brunner NA, Schiffer G, Lampe T, Pohlmann J, Brands M, et al. Identification and characterization of the first class of potent bacterial acetyl-CoA carboxylase inhibitors with antibacterial activity. J Biol Chem. 2004;279: 26066–26073. doi:10.1074/jbc.M402989200

81. Racusen D, Foote M. A major soluble glycoprotein of potato tubers. J Food Biochem. 1980;4: 43–52. doi:10.1111/j.1745-4514.1980.tb00876.x

82. Qiao Y, Piao R, Shi J, Lee S-I, Jiang W, Kim B-K, et al. Fine mapping and candidate gene analysis of *dense and erect panicle 3, DEP3*, which confers high grain yield in rice (*Oryza sativa* L.). Theor Appl Genet. 2011;122: 1439–1449. doi:10.1007/s00122-011-1543-6

83. Hirooka Y, Homma K, Shiraiwa T, Makino Y, Liu T, Xu Z, et al. Yield and growth characteristics of erect panicle type rice (*Oryza sativa* L.) cultivar, Shennong265 under various crop management practices in Western Japan. Plant Prod Sci. 2018;21: 1–7. doi:10.1080/1343943X.2018.1426993

84. Clevenger JP, Korani W, Ozias-Akins P, Jackson S. Haplotype-based genotyping in polyploids. Front Plant Sci. 2018;9. doi:10.3389/fpls.2018.00564

85. Clevenger JP, Ozias-Akins P. SWEEP: A tool for filtering high-quality SNPs in polyploid crops. G3 Bethesda Md. 2015;5: 1797–1803. doi:10.1534/g3.115.019703

